# Unraveling the YAP1-TGFβ1 axis: a key driver of androgen receptor loss in prostate cancer-associated fibroblasts

**DOI:** 10.1101/2025.02.25.640167

**Authors:** Elena Brunner, Elisabeth Damisch, Melanie Emma Groninger, Lukas Nommensen, Lucy Neumann, Georgios Fotakis, Zlatko Trajanoski, Georg Schäfer, Christian Ploner, Sofia Karkampouna, Francesco Bonollo, Marianna Kruithof-de Julio, Natalie Sampson

**Author notes:** Corresponding author: Medical University of Innsbruck, Innrain 52, A6020 Innsbruck, Austria. these authors contributed equally to this manuscript (Elena Brunner, Elisabeth Damisch, Melanie Emma Groninger).

## Abstract

Due to their pivotal roles in tumor progression and therapy resistance, cancer-associated fibroblasts (CAF) are considered key therapeutic targets with loss of stromal androgen receptor (AR) a poorly understood hallmark of aggressive prostate cancer (PCa). A paucity of pre-clinical models however has hampered functional studies of CAF heterogeneity. We demonstrate that our newly-generated CAF biobank contains three FAP^+^-fibroblast subtypes, each with unique molecular and functional traits. Cultures with an early-activated phenotype expressed the highest levels of AR and exhibited AR-dependent growth. Consistently, stromal cells expressing early-activation markers co-expressed nuclear AR in clinical specimens and were enriched in pre-neoplastic lesions/low-grade PCa. Conversely, myofibroblastic CAF (myCAF), which expressed low AR levels *in vitro* and *in vivo* and were proliferatively-insensitive to AR signaling modulation, constituted the predominant CAF subpopulation in stromogenic high-grade PCa and castration-resistant LACP9 patient-derived xenografts. Exacerbation of the myCAF state upon castration of LAPC9-bearing hosts underscored these findings. Mechanistically, AR loss in myCAF was driven by an NFκB-TGFβ1-YAP1 axis, whose combined targeting synergistically repressed myofibroblastic hallmarks and impaired autophagic flux, effects that were potentiated by enzalutamide resulting in myCAF cell death. Collectively, these findings provide a mechanistic rationale for adjuvant targeting of the YAP1-TGFβ signaling axis to improve patient outcomes.

## Introduction

Prostate homeostasis is maintained via extensive stromal-epithelial crosstalk^1^. Perturbation via pathological processes, including prostatic intraepithelial neoplasia (PIN) and prostate cancer (PCa), lead to a stromogenic reaction characterized by deposition of a collagen-rich extracellular matrix (ECM), SMC dedifferentiation and the emergence of cancer-associated fibroblasts (CAF) ^1^. CAF represent a central onco-modulatory component of the tumor microenvironment (TME) that influence PCa progression and therapy response via diverse mechanisms^2^. Since stromal activation occurs early during disease development and positively correlates with parameters of poor prognosis in PCa^3^, there is considerable interest in developing stromal-targeted therapies.

Treatment options for advanced prostate cancer (PCa) are broadly categorized into radiotherapy, taxane-based chemotherapeutics and suppression of the androgen signaling axis via androgen deprivation therapy (ADT) and androgen receptor signaling inhibitors (ARSI) ^4^. While chemotherapy and radiation primarily target highly proliferating cancer cells, ADT and ARSI aim to restrict PCa cells, whose proliferation is dependent on signaling by the androgen receptor (AR), an androgen-responsive transcription factor. Besides luminal epithelial cells, prostatic AR is also expressed by fibroblasts, activated endothelial cells and parenchymal smooth muscle cells (SMC) ^5^, the latter constituting the most abundant stromal cell type of the benign prostate. In PCa however, there is a well-characterized but poorly understood loss of stromal AR, which correlates with poor prognosis^5^. Together with the low stromal proliferative index, current systemic treatments therefore fail to specifically target the stromal component of disease. In stark contrast to the well-characterized pro-proliferative role of epithelial AR, stromal AR is less well understood and appears to exert tumor-suppressive functions^6, 7, 8^. The current lack of prostatic stromal cell models expressing endogenous AR however has limited our understanding of stromal AR function, an issue that must be overcome to clarify the clinical implications regarding potential inadvertent effects of ADT/ARSI on onco-suppressive stromal AR signaling.

CAF are highly heterogeneous and both tumor-suppressive and tumor-promoting subpopulations have been described^2^. The prevailing consensus is that fibroblasts reside on an activation continuum whereby tumor stage and spatial signaling cues play key roles in determining inflammatory (inflammatory CAF, iCAF) and matrix-producing contractile CAF (myofibroblastic CAF, myCAF) phenotypes^9, 10^. Antigen-presenting CAF (apCAF) constitute a third but lesser understood CAF subpopulation that may represent an intermediate iCAF substate^10^. Whilst myCAF populations are universally associated with aggressive disease, functional analyses of CAF subpopulations are largely lacking, primarily due to the lack of appropriate experimental models.

This study aimed to generate an experimentally-versatile fibroblast biobank encompassing fibroblast subpopulations from clinical PCa tissues. The biobank generated contains three distinct primary human prostate FAP^+^ fibroblast subtypes displaying molecular parallels to early-activated fibroblasts (iCAF), those in an intermediate or transitioning phase, and late-activated myCAF substates. High AR expression and sensitivity of the early-activated clinically-favorable fibroblastic state to AR signaling modulation contrasted with the low AR expression of late-activated onco-supportive myCAF, which were proliferatively-insensitive to modulation of AR signaling and androgen deprivation. These findings were supported by studies in clinical specimens and castration-resistant patient-derived xenograft (PDX) models and have potential implications for current ADT strategies. We identified TGFβ1-YAP1 signaling as a central driver of AR loss and acquisition of the late-activated myCAF state. Combined targeting of this signaling axis in myCAF cultures synergistically attenuated their myofibroblastic traits and was associated with hallmarks of impaired autophagy. Enzalutamide potentiated these effects resulting in myCAF cell death. These findings illustrate the translational value of the biobank and suggest that adjuvant targeting of the TGFβ1-YAP1 signaling axis may improve patient response.

## Results

### Generation of a biobank comprising distinct primary prostate fibroblast substates

To better functionally characterize fibroblast heterogeneity within the PCa microenvironment, we developed a biobank of primary fibroblast explant cultures isolated from PCa-enriched and benign-adjacent surgical resections (Supplemental Fig. 1). This biobank currently comprises 396 cultures from 177 biopsy cores of 95 different patients and includes patient-matched cultures from malignant and benign-adjacent regions from 64 different patients (Supplemental Table 1). All cultures tested uniformly expressed mesenchymal markers vimentin and THY1/CD90 but lacked the epithelial marker pan-cytokeratin (Supplemental Fig. 1-2).

Morphological and phenotypic heterogeneity among these cultures prompted us to discern the transcriptomic profile of thirty-eight randomly selected patient-matched cultures (Fig. 1; Supplemental Fig. 3A-B). Bioinformatic analyses identified four distinct clusters (Fig. 1A-C). One cluster (“cycling”) comprising just two samples substantially deviated from the other clusters and was enriched for gene ontology (GO) terms related to cell division (Supplemental Fig. B) similar to previously reported “cycling CAF” ^11^. These samples were not analyzed further. *VIM* was uniformly expressed across the remaining three clusters (C1-C3), however canonical CAF marker expression differed significantly (Fig. 1D and Supplemental Fig. 3C-D).

**Figure 1.**
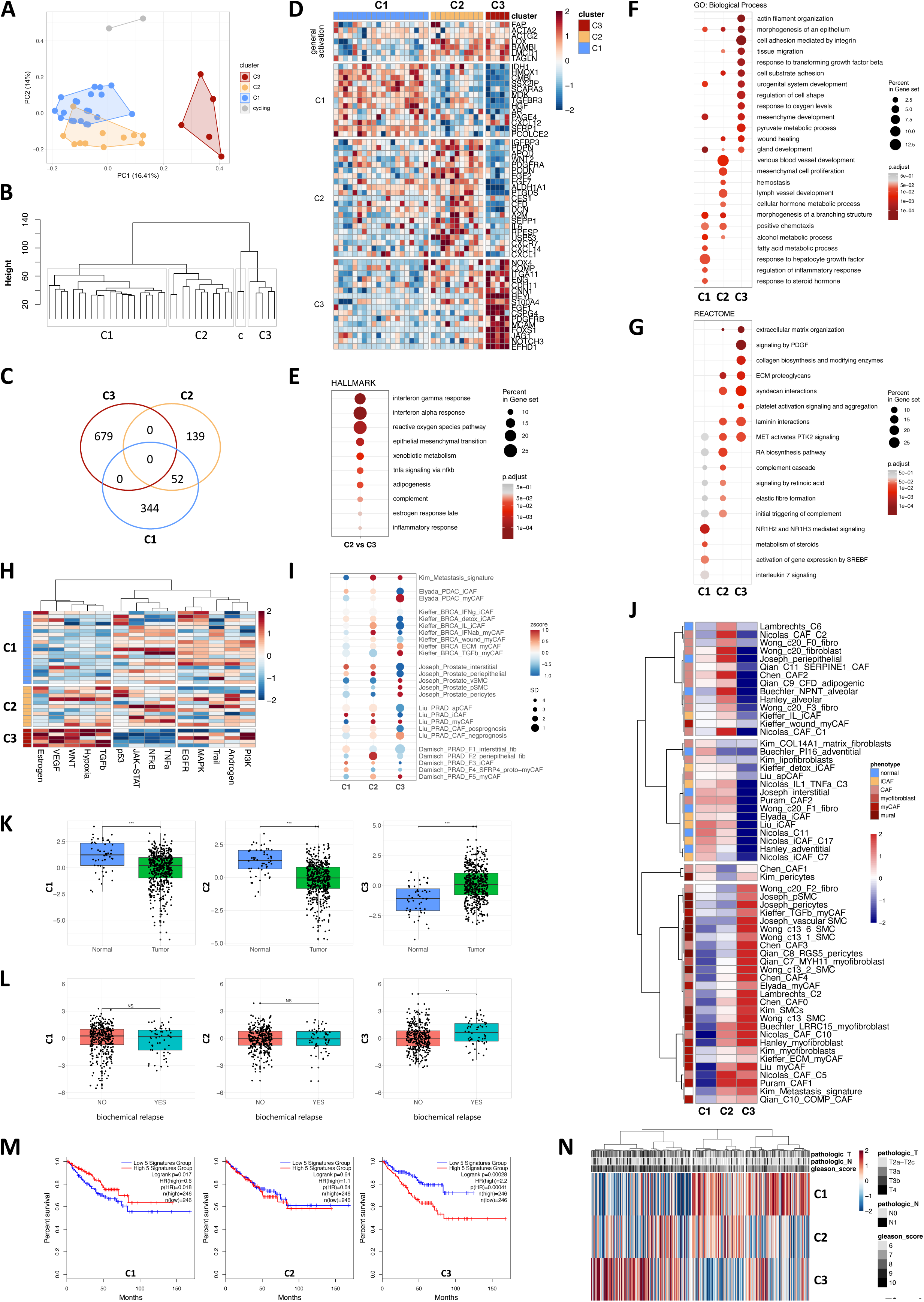
Expression profiling of primary prostate explant cultures identifies distinct fibroblast subpopulations. A-H) Expression profiling of 38 patient-matched primary prostate explant cultures. A) PCA analysis, B) unsupervised hierarchical clustering (c, cycling CAF), C) Venn diagram of significantly upregulated genes and D) sample level expression of selected significantly upregulated genes in each subcluster. E) Hallmark pathways for genes significantly upregulated in C2 *vs.* C3. F) GO Biological Process and G) Reactome pathways for significantly upregulated genes in each subcluster. H) PROGENy inferred signaling pathway activities of each subcluster based on significantly upregulated genes in each subcluster. I) Dotplot comparison and (J) heatmap with rows clustered by correlation to published scRNA-seq datasets using combined z-scores and represented as mean per fibroblast cluster). K-N) Expression of cluster-specific signature genes (combined z-score) for each primary fibroblast cluster in the TCGA-PRAD cohort. N) Heatmap showing combined z-scores of cluster-specific signature genes across all primary tumor samples in the TCGA-PRAD cohort. M) Kaplan-Meier plots using substate-delineating genes foremost annotated to the fibroblast lineage in a PCa scRNA-seq dataset^90^ (Supplemental Fig. 3G) depicting DFS for each signature using median as group cut-off. Source data for panels C, E-L are provided in the Source Data file.

**Figure 2.**
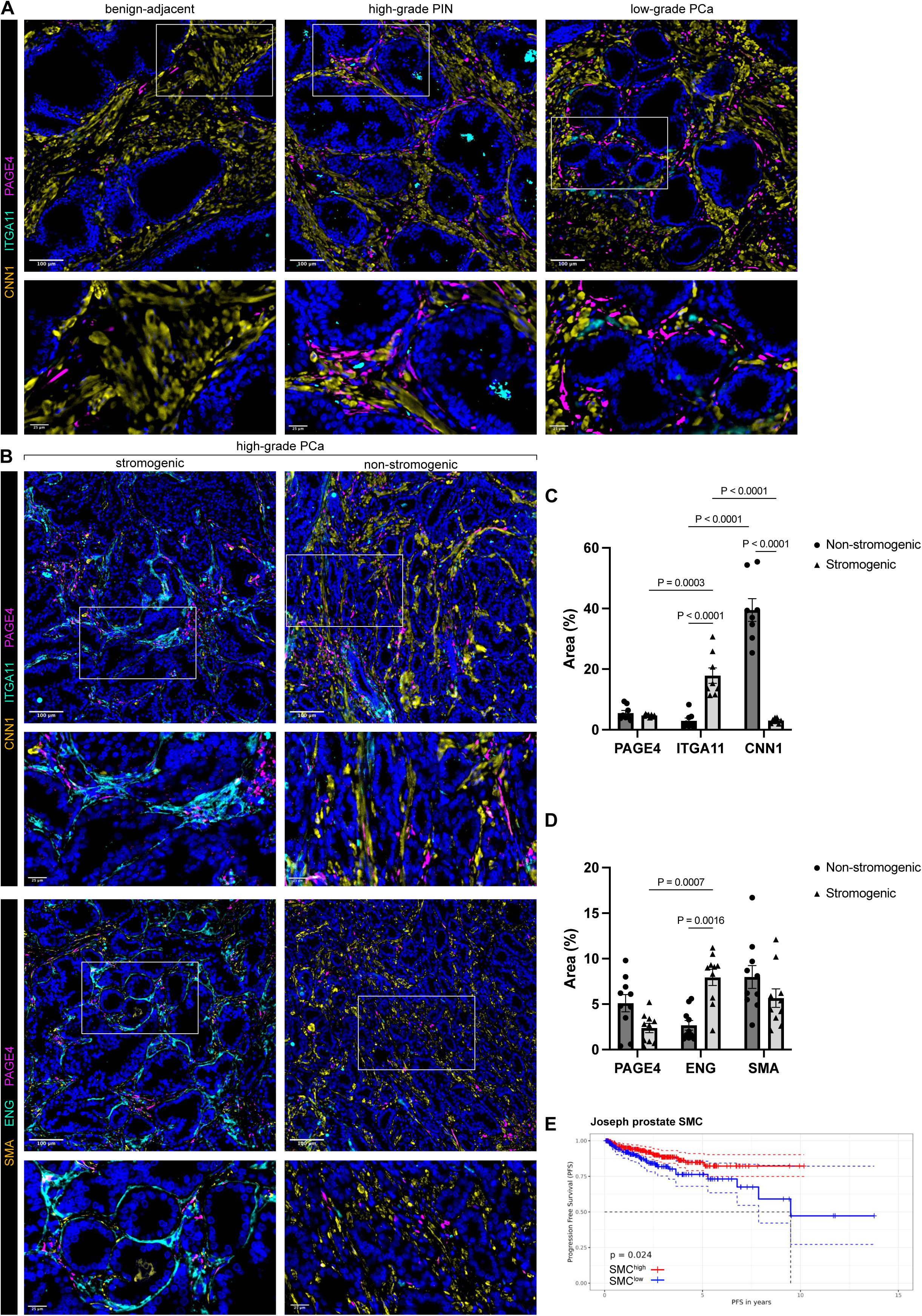
CAF subpopulations are dynamic during prostate cancer progression A-B, D) Human prostate tissue sections of indicated pathology were stained using the antibodies shown. Font color denotes pseudo-coloring in the displayed merged images. Nuclei were counterstained using Hoechst 33342. Boxed regions are shown enlarged beneath each parental image. Images are representative of 4 independent experiments using tissue sections from 8 different patients. C-D) ImageJ quantification of tissues stained as in B) whereby the area stained by the indicated antibody is expressed as a percentage of the total DAPI^+^ area. Data represent mean ± SEM of at least eight fields of view from three different high-grade tumors. E) Kaplan-Meier plots depicting progression-free survival (PFS) of high-grade (≥T stage T2c) samples of the TCGA-PRAD cohort stratified as described in Materials and Methods for high (SMC^high^) or low (SMC^low^) expression levels of the indicated SMC signatures. Source data for panel E is provided in the source data file.

C1 displayed low expression levels of archetypal CAF markers, upregulation of HGF response and MAPK/EGFR signaling, and was enriched for gene sets related to development, morphogenesis and metabolic pathways, (Fig. 1D-G and Supplemental Fig. 3B) suggesting a homeostatic function typically associated with benign prostate fibroblasts. Supportively, C1 exhibited similarity to Buechler *PI16* steady-state “universal” fibroblasts^12^ and Nicolas C11 benign fibroblasts^13^ (Fig. 1I-J, Supplemental Fig. 3E). Similarity to some iCAF-associated signatures however suggested an early activation/pro-inflammatory phenotype that is frequently observed in tumor-adjacent benign tissue^14^. iCAF typically exhibit more tissue-specific differences than myCAF, which are highly similar across different tissues^14^.

Indeed, C1 displayed several prostate-related characteristics, including *AR* expression and upregulation of steroid hormone metabolism and androgen response pathways (Fig. 1D, 1F-H). C2 and C3 both expressed high levels of CAF-associated activation markers (Fig. 1D, Supplemental Fig. 3C-D). C2 however exhibited abundant expression of iCAF-associated markers and was enriched for the iCAF-associated pathways complement activation, IFNγ/IFNα/TNFα pathways as well as signaling via their downstream mediators NFκB and JAK/STAT (Fig. 1D-H; Supplemental Fig. 3B). Indeed, C2 exhibited similarity to several iCAF subsets distinct to those of C1, including Kieffer IL iCAF^15^, Nicolas cluster 3 IL1 TNFa iCAF^13^, Chen early PCa CAF^16^, and *PDPN^+^/SELENOP^+^/SMA^low^*CAF-S5^17, 18^ (Fig. 1I-J, Supplemental Fig. 3E). C2 also displayed weak similarity to some early stage myCAF signatures, including Kieffer IFNαβ-myCAF but not late stage TGFβ- or ECM-myCAF^10, 15^, suggesting initial transitioning on the myCAF trajectory (Fig. 1I-J and Supplemental Fig. 3E).

C3 on the other hand displayed significant upregulation of canonical myCAF genes *ITGA11, SPP1, CNN1* and *ENG* and was enriched for hallmark myCAF pathways, such as ECM organization, collagen biosynthesis, cell adhesion mediated by integrin, glycolysis, hypoxia, TGFβ and VEGF pathways (Fig. 1D-H; Supplemental Fig. 3B) in keeping with acquisition of contractile/myofibroblastic properties. Consistently, C3 demonstrated strong similarity to myCAF signatures from diverse cancer types, including breast, prostate, pancreas and lung cancers (Fig. 1I-J).

Late activation/myCAF phenotypes positively correlate with hallmarks of poor prognosis across multiple tumor types^19^, whereas early activation/iCAF phenotypes are frequently associated with positive clinicopathological parameters^14^. Concordantly, C3 marker genes were significantly elevated in the TCGA-prostate adenocarcinoma (PRAD) cohort compared to benign controls and positively correlated with poor prognostic indicators, including biochemical relapse (Fig. 1K-N; Supplemental Fig. 3F). In contrast, C1 and C2 marker genes were significantly lower in primary PCa relative to normal controls with the C1 signature positively associated with increased disease-free survival (DFS; hazard ratio (HR) 0.6; *P*=0.018). In contrast, the C3 signature was associated with decreased DFS (HR 2.2; *P*=4.1×10^-4^) (Fig. 1M).

The presence of distinct fibroblast substates within our biobank was validated via quantitative real time PCR (qRT-PCR) of fibroblast activation and substate-delineating markers in an independent cohort of twenty-nine explant cultures as well as in three cultures from the original bulk transcriptomic dataset (qRT-PCR cohort, Supplemental Fig. 4A). Expression levels of C1, C2 and C3 markers displayed longitudinal stability within an individual culture (Supplemental Fig. 4B). Immunofluorescent staining for the C2 markers PDGFRA and PDPN and C3 marker ITGA11 readily distinguished cultures in line with their expression profile-based hierarchical classification (Fig. 1A, 2A, 2C, Supplemental Fig. 4C). We also performed multiparametric flow cytometry (FC) using a panel of seven stromal markers for thirty-six cultures, including ten samples from the qRT-PCR cohort and four samples from the transcriptomic cohort (Supplemental Fig. 5). All samples were positive for FAP reminiscent of the heterogeneous FAP^+^ CAF-S1 superpopulation from breast cancer^15, 20^. Dimensional reduction and unsupervised clustering based on prominently expressed stromal markers initially identified thirteen clusters (Supplemental Fig. 5A-C). Gating for these clusters based on their corresponding cell surface expression levels revealed eight main clusters (Supplemental Fig. 5D-E). Notably, cultures annotated to a particular fibroblast subpopulation in the transcriptomic and qRT-PCR cohorts displayed high concordance with their grouping in the FC dataset with each fibroblast subpopulation demarcated by a distinct gated FC cluster (Supplemental Fig. 5F-H). Early-activated fibroblasts (C1) and transitioning CAF (C2) could be differentiated from C3 myCAF via PDPN consistent with multiple studies demonstrating *PDPN* expression in iCAF^17, 18, 21, 22, 23^. In turn, PDPN^+^ C1 and C2 cultures were distinguishable via their differential cell surface expression of MCAM. The C3-enriched cluster 1 was demarcated by lower PDPN and PDGFRA expression but moderate expression of MCAM in accordance with previously reported myCAF^11, 24^. Fluorescence-activated cell sorting (FACS) of cultures for clusters 6, 10 or 1 and subsequent qRT-PCR confirmed high expression levels of C1, C2 or C3-delineating markers in cells sorted for FC gated clusters 6, 10 and 1, respectively (Supplemental Fig. 5I). In summary, the fibroblast biobank encompasses phenotypically-stable cultures that display extensive molecular similarities to early (C1), intermediate (C2) and late (C3) fibroblast activation states that have been identified *in situ* across multiple cancer types.

### Explant cultures display molecular parallels to distinct CAF substates in clinical PCa

We next aimed to validate key markers that delineated C1-C3 explant cultures in clinical PCa. To this end, patient tissues were immunostained for the SMC markers CNN1, CCDC102B or SMA (encoded by *ACTA2*), the C1 marker PAGE4, C2 marker CES1 and C3 markers ITGA11 or ENG (Fig. 2 and Supplemental Fig. 6). ITGA11, CES1 and PAGE4 were sparsely expressed in benign-adjacent tissues, which was predominated by SMC (Fig. 2A and Supplemental Fig. 6A-C) with ENG expression restricted to the endothelium of some vessels as previously reported^25^. PAGE4 and CES1 however were markedly upregulated high-grade PIN (HGPIN) and low-grade PCa being localized to both the periglandular and interstitial stroma. In contrast, ITGA11 and ENG expression remained very low in HGPIN but an increased frequency of cells expressing low/moderate levels of ITGA11 or ENG was observed in low-grade PCa. Across each of these stages, SMC remained the predominant parenchymal stromal cell type.

Two distinct types of high-grade tumors were identified: those with a considerable proportion of parenchymal SMC yet low numbers of ITGA11^+^/ENG^+^ CAF *vs.* tumors with a near complete loss of parenchymal immunoreactivity for the mature SMC differentiation marker CNN1 and lower levels of the general SMC marker SMA but a high abundance of ITGA11^+^/ENG^+^ cells that were predominantly restricted to the peri-tumoral space (Fig. 2B-D). These distinct stromal compositions are consistent with previously described “non-stromogenic” and “stromogenic” PCa, respectively^3^. Whilst the abundance of PAGE4^+^ or CES1^+^ cells did not significantly differ between these two tumor types, PAGE4^+^/CES1^+^ cells tended to accumulate in nests within intraglandular areas of stromogenic tumors. This pattern was not readily apparent in non-stromogenic high-grade tumors (Fig. 2B and Supplemental Fig. 6A) suggesting the presence of defined spatial CAF niches in stromogenic advanced PCa. In further support of the clinical relevance of ITGA11^+^/ENG^+^ stromogenic *vs.* non-stromogenic tumors, we observed that progression-free survival (PFS) of patients with pathological tumor stage ≥T2c was significantly higher for those with tumors expressing high levels of SMC markers compared to those with lower expression of SMC markers (Fig. 2E, Supplemental Fig. 6H-I). In summary, PCa progression is associated with dynamic changes in CAF subpopulations whereby the C3 markers ENG and ITGA11 demarcate the predominant CAF substate in stromogenic high-grade tumors with potential implications for improved prognostic staging.

### CAF subpopulations display distinct phenotypic hallmarks

A paucity of *in vitro* models has limited our understanding of CAF heterogeneity at the functional level. We therefore employed C1 and patient-matched C2 or C3 explant cultures representing early-activated (C1), intermediate (C2) and late activation (C3) fibroblast states from approximately thirty patients for an initial functional characterization. C1 cultures displayed a typical spindle-like, light-refractive fibroblast morphology, whereas both C2 and C3 cultures exhibited greater 2-dimensional (2D) spreading with suspensions of C3 cells slightly but significantly larger than C2 and patient-matched C1 cells (Fig. 3A-B). No differences were observed in cell viability. However, C2 and C3 explant cultures proliferated slower than patient-matched C1 (Fig. 3C-D). Consistent with their myCAF annotation, C3 cultures displayed significantly higher migration and collagen-1 gel contraction than either C2 or patient-matched C1 (Fig. 3E-H). Since CAF-led migration promotes cancer cell invasion and migration^26^, this finding further implicates the late/myCAF phenotype in promoting aggressive PCa.

**Figure 3.**
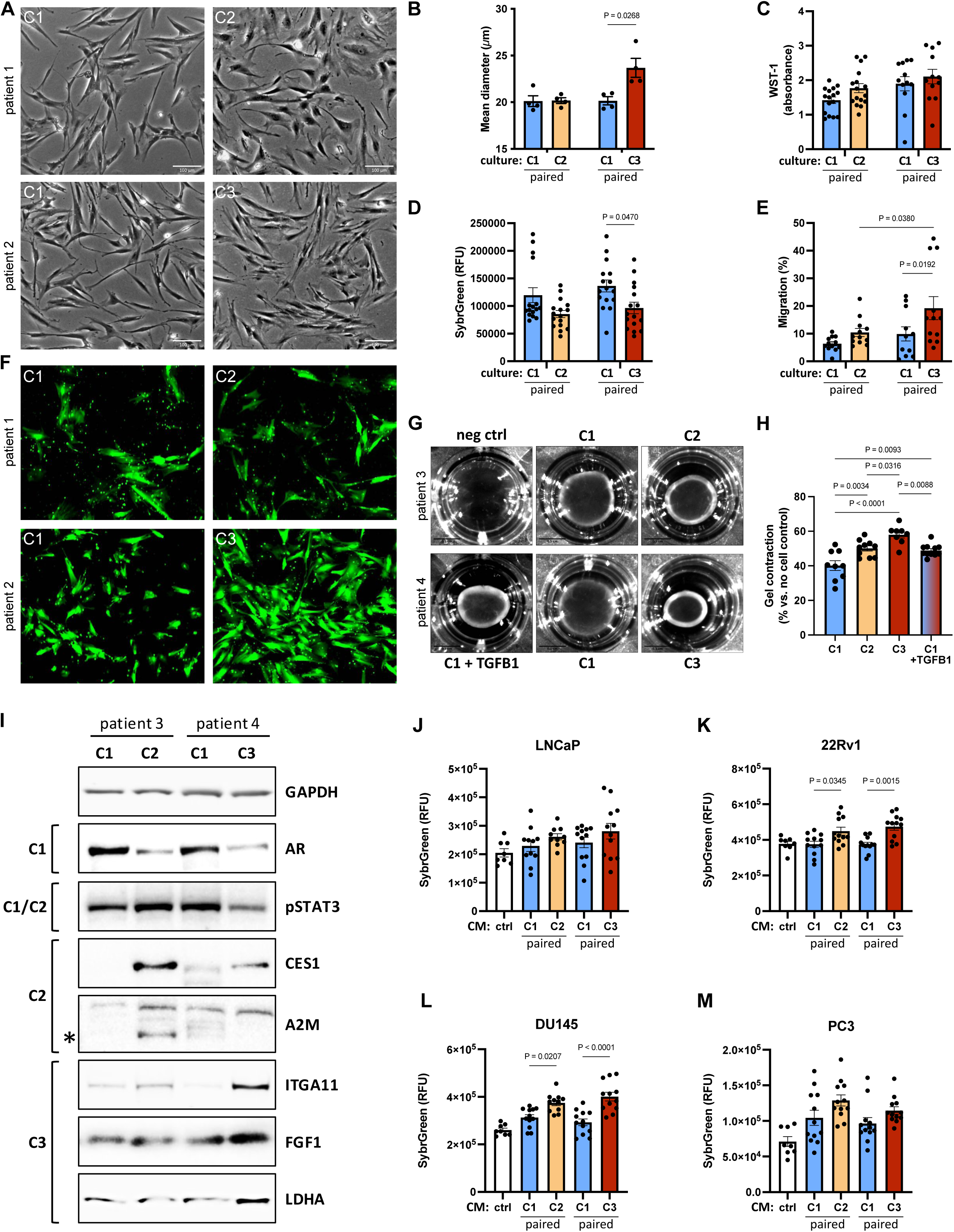
Primary CAF cultures exhibit distinct functional hallmarks yet shared onco-supportive capacities Explant cultures displaying a C2 or C3 molecular profile and patient-matched C1 cultures were characterized via A) brightfield imaging, B) cell size, C) cell viability via WST-1 assay, D) proliferation via SybrGreen staining, E-F) transwell migration, G-H) collagen I gel contractility and I) Western blotting with the antibodies indicated (* denotes the A2M band). J-M) Proliferation of J-K) AR^+^ or L-M) AR^-^ PCa cell lines incubated with CM from C2 or C3 cultures and patient-matched C1s. Non-conditioned media (nonCM) incubated for the same duration without primary fibroblasts served as control. Data represent mean fluorescence of quadruplicate wells from independent experiments using fibroblasts derived from three different donors. Statistical significance was calculated: B-E) two-way ANOVA with Holm-Šídák multiple comparison correction, H, K) one-way ANOVA with Tukey multiple comparison correction. A, F, G, I) images are representative of at least three different experiments using primary cells isolated from different donors. Ctrl, control.

We next sought to validate at the protein level key components of pathways enriched in each of the culture subpopulations (Fig. 3I). C2 displayed the strongest expression of carboxylesterase 1 (CES1), an NFκB-regulated alternatively-spliced serine hydrolase involved in lipid/cholesterol metabolism^27^, a finding consistent with their upregulation of adipogenesis (Fig. 1E, Supplemental Table 9). Likewise, C2 cultures expressed alpha-2-macroglobulin (A2M), which acts as an intra- and extra-cellular chaperone of inflammatory cytokines, including IL1β, and activates the complement cascade^28^, an iCAF hallmark pathway that was also upregulated in C2 cultures (Fig. 1G). Pro-inflammatory cytokines induce iCAF formation at least in part via JAK/STAT signaling^29^. *In silico* analyses implicated TNFα and interferons gamma and alpha and their downstream signaling mediator STAT3 in C2 cultures (Fig. 1E, 1H, Supplemental Table 9; Supplemental Fig. 3B). Accordingly, C1 and C2 cultures displayed abundant levels of STAT3 phosphorylated at tyrosine 705 (pSTAT3) compared to C3 cultures (Fig. 3I). Consistent with enrichment of steroid hormone metabolism/signaling (Fig. 1F-H), elevated AR expression in C1 cultures was also confirmed at the protein level. C3 cultures on the other hand exhibited low levels of AR but high levels of FGF1 and the TGFβ-responsive and collagen type I-binding integrin ITGA11 (Fig. 3I). The “reverse Warburg effect” is a key myCAF hallmark^30^. Accordingly, C3 cultures displayed the highest levels of LDHA, indicating a strong glycolytic activity. Supportively, the lactate transporter MCT4 and HIF1A, which are involved in promoting glycolysis and myofibroblast differentiation^30^, were significantly upregulated (Supplemental Table 3), suggesting C3 cultures may be highly active in supporting cancer cell metabolism and growth.

CAF can either support or restrict cancer growth by secreting pro-inflammatory and mitogenic growth factors^1^. We therefore assessed proliferation of PCa cells following exposure to conditioned media (CM) from C1, C2 or C3 cultures or non-conditioned media as control. When exposed to CM from C1 cultures, PCa cells proliferated at similar rates to control media, regardless of their AR status, indicating that C1 cultures do not significantly influence cancer cell growth. Conversely, CM from C3 cultures, which are linked to poor clinical outcomes, significantly increased the proliferation of both AR-positive and AR-negative PCa cell lines (Fig. 3J-M). This suggests that C3 cultures strongly promote cancer cell growth. A similar effect was observed with CM from C2 cultures. Collectively, these studies indicate that the different fibroblast substates are not only molecularly divergent but also have varying effects on cancer cell growth, with some promoting proliferation more than others.

### AR is downregulated in myofibroblastic CAF

Loss of stromal AR in PCa is an established but poorly understood indicator of poor prognosis^31^. Since C1 cultures expressed higher levels of AR than C3 cultures and were enriched for GO terms/hallmark pathways related to androgen response and metabolism (Fig. 1D, 1F-H; Supplemental Fig. 3B, Fig. 3I, Fig. 4A), subsequent studies focused on exploiting our biobank to investigate endogenous AR function in primary prostate fibroblasts in the context of PCa. First, we confirmed elevated expression of *AR* in C1 and its downregulation in C3 cultures via qRT-PCR in samples remaining from the independent cohort (Fig. 4B) as well as at the protein level in additional biobank cultures (Fig. 4C and Supplemental Fig. 7A). AR protein levels exhibited a non-significant trend towards inverse correlation with the C3 marker ITGA11 but significantly positively correlated with pSTAT3 levels (Fig. 4C-D, Supplemental Fig. 7A-B), a hallmark of C1/C2 cultures (Fig. 1H) and key pathway of the iCAF/early CAF phenotype^32^.

**Figure 4.**
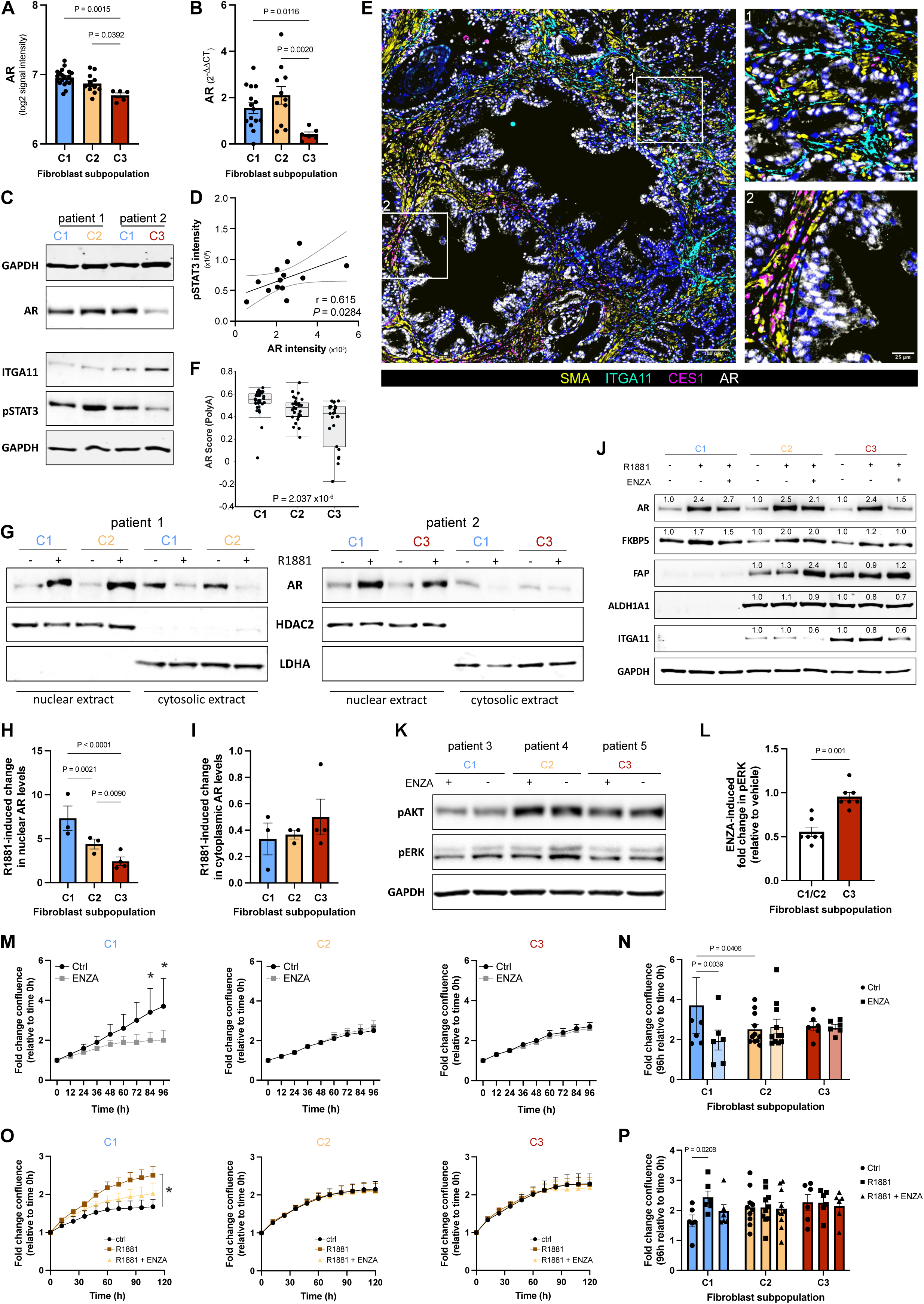
AR loss in myofibroblastic CAF is associated with proliferative-insensitivity to enzalutamide AR expression levels A) in explant cultures of the original transcriptomic dataset and B) via qRT-PCR in the independent explant cohort and C) via Western blotting. D) Densitometric-based correlation of AR and pSTAT3 protein levels in thirteen explant cultures. E) Immunofluorescent staining of a human prostate tissue section comprising prostate cancer and benign-adjacent tissue using the antibodies indicated. Font color denotes pseudo-coloring in the displayed merged images. Nuclei were counterstained using Hoechst 33342. Boxed regions are shown enlarged, right. F) AR score of samples from the SU2C metastatic PCa cohort grouped according to expression of fibroblast substate-delineating markers. G) Western blotting and H-I) densitometric quantification of AR in nuclear and cytosolic extracts incubated with or without 10 nM R1881 for 1h. J) Western blotting using the antibodies indicated of fibroblast cultures pretreated with 10 µM enzalutamide (ENZA) or vehicle equivalent before addition of 10 nM R1881 for a further 72 h. Values denote normalized fold change in protein levels relative to the corresponding control. K) Western blotting using the antibodies indicated and L) densitometric quantification of fibroblast cultures incubated for 72h with 10 µM enzalutamide (ENZA) or vehicle equivalent. M-P) Real time proliferation of the indicated fibroblast culture incubated M-N) with 10 µM enzalutamide or vehicle equivalent (ctrl) or O-P) 1 pM R1881 with 10 µM enzalutamide or vehicle equivalent. N, P) Bar chart denoting proliferation of cultures after 96h from M) or O) respectively. Statistical significance was calculated: A-B, H-I) one-way ANOVA with two-stage step-up method of Benjamini, Krieger and Yekutieli multiple comparison correction; D) Pearson correlation; F) Kruskal-Wallis test; L) Mann-Whitney test; M-P) two-way ANOVA with Holm-Šídák multiple comparison correction. C, E, G, J-K) images are representative of at least three independent experiments using primary material from different donors.

As previously reported, stromal AR was primarily expressed in parenchymal SMC of the benign prostate but decreased in PCa (Supplemental Fig. 7C). We noted however nuclear AR expression in stromal cells negative for SMC markers that were interspersed between tumorigenic glands, strongly reminiscent of the aforementioned spatial distribution of PAGE4^+^/CES1^+^ CAF (Fig. 2, Supplemental Fig. 7C). Indeed, multiplex staining revealed co-expression of nuclear AR with the early/intermediate C1-C2 markers CES1 and PAGE4 (Fig. 4E and Supplemental Fig. 7D-F). In contrast and consistent with the above *ex vivo* findings, AR was expressed at significantly lower levels in ITGA11^+^/ENG^+^ myCAF in clinical PCa and was significantly decreased in samples of the SU2C metastatic cohort exhibiting a C3 profile (Fig. 4E-F and Supplemental Fig. 7D-F). These data raised the possibility that by virtue of their differential AR expression, distinct fibroblast substates may be differentially sensitive to androgens and ADT/ARSI, such as enzalutamide.

### AR downregulation in myCAF is associated with insensitivity to the anti-proliferative effects of enzalutamide

We thus next sought to assess AR functionality among the distinct fibroblast substates encompassed within our biobank (Fig. 4G-P). Despite different total AR levels (Fig. 4A-C), the synthetic androgen R1881 rapidly induced AR nuclear translocation in cells pre-incubated under steroid hormone-depleted conditions, regardless of the fibroblast substate (Fig. 4G). In C3 cultures however, ligand-induced nuclear translocation was significantly lower than that induced in C1 or C2 cultures (Fig. 4H). Concordantly, the corresponding decrease in cytosolic AR levels in response to R1881 was markedly attenuated in C3 cultures compared to C1/C2 (Fig. 4I). Target genes modulated by endogenous stromal AR remain poorly defined. However, R1881 treatment of C1/C2 cultures modestly increased and decreased the previously reported positively and negatively AR-regulated genes FKBP5 and FAP, respectively^7^ in an enzalutamide-sensitive manner at both the mRNA and protein level (Fig. 4J and Supplemental Fig. 7G-H). No discernible change however was observed in parallel-treated C3 cultures despite R1881-mediated increased total AR protein levels across all substates (Fig. 4J), an observation consistent with ligand-induced post-translational AR stabilization^33^. Similar findings were observed following AR inhibition under steroid hormone-replete conditions for FAP whilst there was no apparent change in FKBP5 levels (Supplemental Fig. 7H). With the exception of a modest but consistent decrease in ITGA11 protein levels that was further exacerbated by enzalutamide, no discernible change was observed in the expression of fibroblast substate markers either at the mRNA or protein level in response to picomolar or nanomolar concentrations of exogenous androgen or upon enzalutamide-mediated AR inhibition (Fig. 4J, Supplemental Fig. 7H-I and data not shown).

AR also engages in crosstalk with intracellular signaling pathways. Indeed, whilst enzalutamide treatment under steroid hormone-replete conditions did not alter phospho-AKT levels, AR inhibition significantly decreased phospho-levels of ERK1/2 in C1 and C2 fibroblasts but not in C3 cultures (Fig. 4K-L and Supplemental Fig. 7J). Furthermore, enzalutamide significantly attenuated the proliferation of C1 cultures, an effect that was not observed in C2 or C3 cultures potentially due to high basal pAKT activity (Fig. 4M-N and Supplemental Fig. 7K), a key regulator of CAF proliferation^34^. Similarly, under steroid hormone-depleted conditions exogenous androgen promoted the proliferation but suppressed the migration of C1 cultures in an AR-dependent manner, whereas this effect was not observed in either C2 or C3 cultures (Fig. 4O-P and Supplemental Fig. 7L-O). Collectively, the aforementioned data indicate that variability in AR levels among the fibroblast substates translates into functional differences with regards to their proliferative sensitivity to modulation of AR signaling.

### The myCAF phenotype is associated with castration-resistance in vivo

Data thus far implicated a potential association of AR^low^ myCAF with castration-resistance. We therefore investigated CAF heterogeneity in the BM18 androgen-sensitive PCa PDX model, which regresses upon host castration, and the castration-insensitive LAPC9 PCa PDX model (Fig. 5A) ^35^. Stromal expression levels of mouse orthologs to C1, C2 and C3 markers were discerned by exploiting reports that infiltrating host stromal cells replace human-derived stroma in serially-passaged PDXs^36^. Whilst general activation and C3 markers were expressed in the intact BM18 model (Fig. 5B and Supplemental Fig. 8A), their expression levels were markedly higher in intact LACP9 xenografts (Fig. 5B and Supplemental Fig. 8A), an observation consistent with the more aggressive phenotype of the LAPC9 model. Furthermore, C1 markers were more abundantly expressed in intact BM18 tumors compared to intact LAPC9 tumors (Fig. 5B and Supplemental Fig. 8A). These findings were confirmed at the protein level for Pdgfra and Itga11 (Fig. 5D-E and Supplemental Fig. 8B-E). Accordingly, an abundance of Itga11^+^ cells was detected throughout the intra-tumoral stroma of intact LAPC9 xenografts (Fig. 5D and Supplemental Fig. 8B-E). Conversely, Itga11 immunoreactivity was restricted to the periphery of intact BM18 tumors with no Itga11^+^ cells detectable in the intra-tumoral stroma (Fig. 5D and Supplemental Fig. 8D).

**Figure 5.**
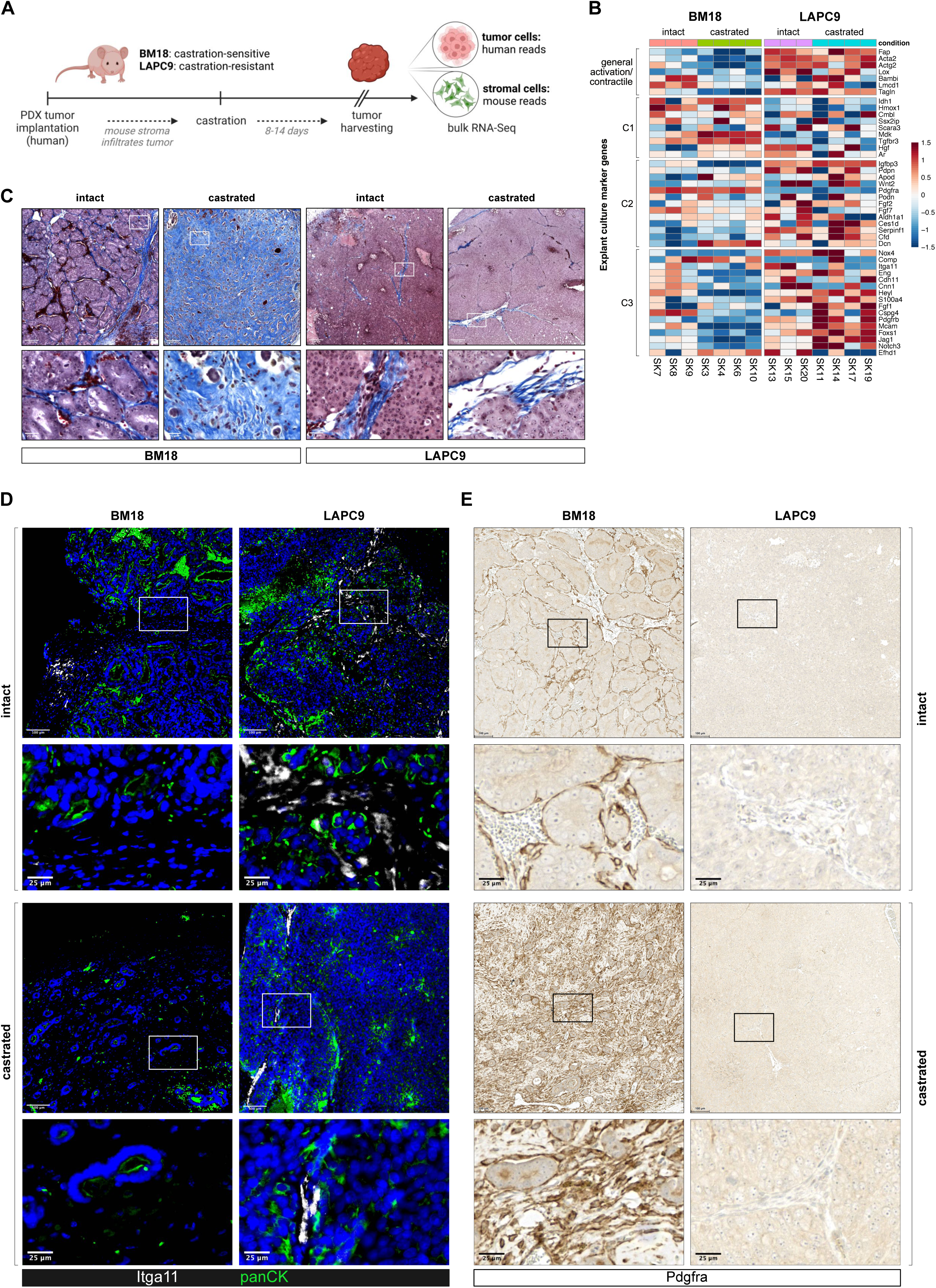
Myofibroblastic CAF are associated with castration-resistance *in vivo* A) Graphical overview of the PDX system whereby host stromal cells infiltrate human PCa PDX tissue replacing human stromal tissue upon serial passaging ^36^. Mouse- and human-specific sequence reads thus represent the stromal and tumoral transcriptomes, respectively. B) Sample level heatmap of murine orthologs of the indicated markers in intact or castrated mice harboring the indicated PDX tumor. C) Masson’s trichrome, D) immunofluorescent and E) immunohistochemical staining of the indicated PDX tumors (*n*=3 tumors stained in four independent experiments). Boxed regions are shown enlarged beneath each parental image. D) Font color denotes pseudo-coloring in the displayed merged images. Nuclei were counterstained using Hoechst 33342. A) Adapted from ^35^.

Notably, upon castration C3 markers were further upregulated in the LAPC9 stroma (Fig. 5B and Supplemental Fig. 8A). Due to the continued exponential tumor cell growth of this castration-resistant model, only a limited amount of stromal tissue remained in the tumors harvested at experimental endpoint. Nonetheless, intra-tumoral Itga11^+^ cells were readily detectable at both the tumor periphery and within the tumor core (Fig. 5D and Supplemental Fig. 8B-D), indicating their insensitivity to androgen deprivation. In contrast, castration-induced regression of BM18 tumors was accompanied by decreased mRNA levels of 14 out of 16 C3 markers and absence of Itga11 immunoreactivity but a concomitant increase in C1 and some C2 markers (Fig. 5B and 5D-E). Collectively, these data implicate the C3/late myCAF substate in castration-resistant PCa and that androgen depletion may favor survival of the clinically-unfavorable AR^low^ myCAF state. Exploring the signaling pathways that drive the C3 substate may provide more actionable targets for therapy.

### TGFβ signaling drives AR loss in C1 and C2 but not C3 cultures

In view of the abovementioned findings, we next sought to delineate intracellular signaling pathways that determine CAF substate identity and drive AR loss. In line with their myCAF phenotype, PROGENy revealed enrichment of TGFβ signaling in C3 cultures (Fig. 1H). Indeed, treatment of C1 cultures with recombinant TGFβ1 significantly upregulated C3 markers whereas C1 and C2 markers, including AR, were concomitantly significantly downregulated (Fig. 6A-B). Enzalutamide treatment neither alone nor in combination with TGFβ1 significantly altered C1, C2 or C3 marker expression.

**Figure 6.**
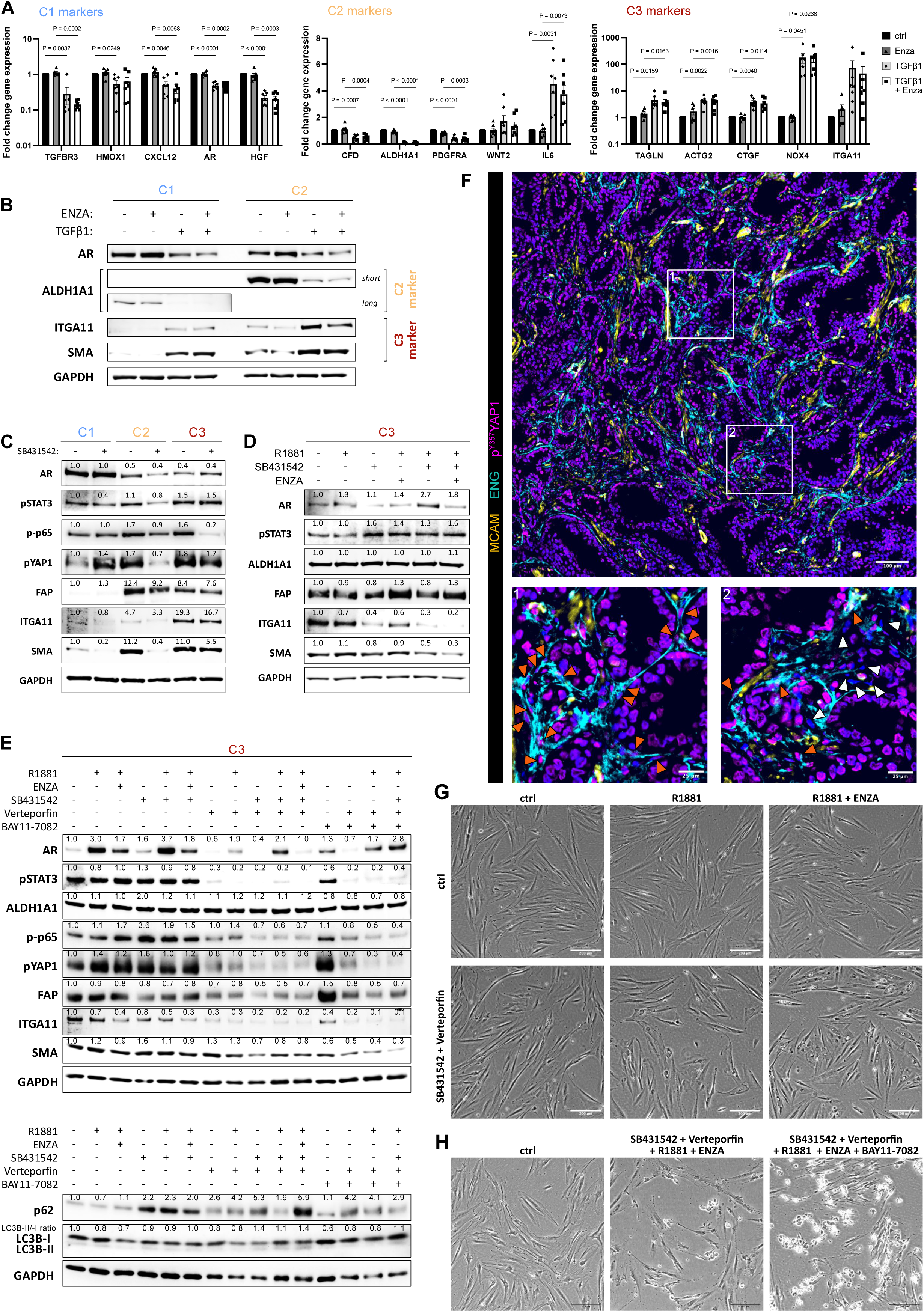
NFκB-YAP-TGFβ signaling axis underlies AR loss during fibroblast phenotypic switching A) qRT-PCR of C1 cultures treated with 10 µM enzalutamide (ENZA) or vehicle control in the presence or absence of 2 ng/ml TGFβ1 for 72h. B) Western blotting of C1 or C2 cultures treated as in A). For ALDH1A1 the membrane was exposed for short or longer durations for optimal signal visualization. C-. E) Western blotting of fibroblast cultures treated as indicated with 1 µM SB431542, 10 nM R1881, 1 µM verteporfin, 2 µM BAY11-7082, 10 µM ENZA or vehicle equivalent for 96h under steroid hormone B) -replete or D-E) -depleted conditions. Values denote normalized fold change in protein levels relative to lane 1. F) Immunofluorescent staining of human prostate cancer tissue using the antibodies indicated. Font color denotes pseudo-coloring in the displayed merged images. Nuclei were counterstained using Hoechst 33342. Boxed regions are shown enlarged below. Orange arrowheads (panels 1 and 2) highlight myCAF nuclei positive for p^Y357^YAP1, pink arrowheads (panel 2) demarcate MCAM^-^ENG^-^ stromal cells expressing low/non-detectable levels of p^Y357^YAP. G) Morphology of fibroblast cultures treated as in E). Statistical significance was calculated: A) two-way ANOVA with two-stage step-up method of Benjamini, Krieger and Yekutieli multiple testing correction. B-G) images are representative of at least three independent experiments using primary material from different donors.

To ascertain whether TGFβ signaling alone determines the fibroblast phenotypic substate, we treated C1, C2 and C3 cultures under steroid hormone-replete conditions with the ALK5 inhibitor SB431542 (Fig. 6C, Supplemental Fig. 9A-B). In line with their low activation profile (Fig. 1 and as further exemplified by their low FAP levels), TGFβ receptor inhibition in C1 cultures yielded no significant changes at the mRNA level. At the protein level, the already low levels of SMA were decreased, suggesting these cultures have a low level of TGFβ signaling, whose attenuation further supports the AR^+^/SMA^low^ C1 phenotype. In contrast, SB431542 treatment of C2 cultures resulted in marked decreased levels of SMA whereby residual ITGA11 levels also modestly decreased implying that the weak myofibroblastic features of C2 cultures are at least in part due to intrinsic TGFβ signaling. Conversely, mRNA and protein levels of C1, C2, C3 or contractile/general activation markers were not significantly modulated upon SB431542 treatment of C3 cultures (Fig. 6C-E and Supplemental Fig. 9A-B), indicating that pathways other than TGFβ are primarily responsible for maintaining the C3 phenotype. Since stromal TGFβ and AR signaling axes engage in suppressive crosstalk^37^, we examined whether R1881 synergizes with TGFβ receptor inhibition to modulate the C3 phenotype. Indeed, in addition to the aforementioned ligand-dependent AR protein stabilization (Fig. 4J), exogenous R1881 under otherwise steroid hormone-depleted conditions further decreased ITGA11 and SMA protein levels compared to either agent alone in a manner intriguingly potentiated by enzalutamide (lanes 5, 6 Fig. 4J and 6D-E).

### A YAP1-TGFβ axis drives AR loss in C3 cultures and maintains the myCAF phenotype

Besides TGFβ signaling, the mechanotransducer Yes-associated protein 1 (YAP1) is also implicated in driving myCAF conversion^38^. Accordingly, compared to C1 cultures basal levels of YAP1 phosphorylated at the activating residue tyrosine 357^39^ (hereon pYAP1) were higher in C2 and C3 cultures and in the latter, were insensitive to TGFβ receptor inhibition (Fig. 6C). Similar levels of pYAP1 in C2 and C3 cultures under basal steroid hormone-replete conditions seems contradictory in view of their significantly different myCAF profiles. However, YAP1 phosphorylation at tyrosine 357 differentially redistributes YAP1 promoter occupancy in a context-dependent manner due to association of YAP1 (that is itself unable to bind DNA) with different transcription factors^40, 41, 42^. Supportively, C2 and C3 cultures displayed differential upregulation of canonical YAP-target genes (Supplemental Fig. 9G) with C3 cultures expressing significantly higher levels of the well-established YAP1-binding partner and transcription factor *VGLL4* compared to C2 cultures (log2FC 0.7; padj 0.02). Furthermore, ENG^+^ cells in stromogenic high-grade PCa displayed abundant nuclear pYAP1 whereas ENG^-^MCAM^-^ stromal cells in the intra-glandular tumoral space primarily associated with early/intermediate CAF (Fig. 2) were largely devoid of nuclear pYAP1 (Fig. 6F).

Compared to SB431542 treatment alone, single agent treatment with the YAP1 inhibitor verteporfin decreased pYAP1, ITGA11, AR and pSTAT3 levels but had no effect on SMA protein levels. Combined treatment of C3 cultures however strongly depleted FAP, SMA and ITGA11 levels (lanes 4, 7, 9 Fig. 6E) with treated cells displaying a more elongated and light-refractive morphology characteristic of C1 cultures (Fig. 6G and Supplemental Fig. 9H). Mechanical changes in cell shape and substrate stiffness potently activate YAP1^43^. Consistently, a similar downregulation of C3 markers and the YAP target genes *CYR61* and *CTGF* was observed when C3 but not C1/C2 cells were cultured as 3D spheroids or on soft (2 kPa) hydrogels as an alternative means to inhibit YAP1 signaling and further implicate its central role in maintaining the C3 phenotype (Supplemental Fig. 9C-F).

Despite C3 marker downregulation and morphological similarity to early-activated cultures, neither C1 nor C2 markers were discernibly upregulated at either the mRNA or protein level upon verteporfin/SB431542 co-treatment, upon culture as 3D spheroids or on soft hydrogels as an alternative means to reduce YAP1 activity. Thus, abrogation of myCAF hallmarks upon TGFβ1-YAP1 targeting does not revert C3 myCAF to an C1 or C2 state.

### Combined YAP1-TGFβ targeting renders deactivated myCAF sensitive to enzalutamide-induced cell death

On soft hydrogels, as 3D spheroids and 2D verteporfin mono-treated cells we noted significant upregulation of the NFκB-regulated pro-inflammatory cytokines *IL6, IL8* and *IL1B* in a manner attenuated by the IKK inhibitor BAY11-7082 (Supplemental Fig. 8D-F and data not shown). Additionally, verteporfin mono-treated C3 cultures exhibited low levels of vacuole formation (Supplemental Fig. 9H-I). Combined TGFβ-YAP1 targeting in the presence of R1881 further increased vacuole formation and interleukin expression levels, suggestive of autophagy. Again, these effects were potentiated by enzalutamide. Cultures from multiple donors responded similarly, however we observed patient-specific differences in the severity of response with some cultures undergoing considerable cell death (Fig. 6G-H, Supplemental Fig. 9H-J and data not shown).

Consistent with its role in recycling proteins and organelles during cellular reprogramming, CAF activation is dependent on autophagy with YAP transcriptional activity required for the generation of autolysosomes from autophagosomes^44^. Defective autophagy leads to the accumulation of the selective autophagy receptor p62, which also acts as a signaling hub to activate pro-survival NFκB signaling^45^. Accordingly, p62 levels robustly increased upon mono-treatment with SB431542 or verteporfin, were further increased upon combined TGFβ-YAP1 targeting, again in a manner potentiated by R1881 and enzalutamide. The ratio of the autophagosomal marker LC3B-II to LC3B-I however was only modestly increased under combined TGFβ-YAP1 targeting in the presence and absence of enzalutamide (Fig. 6E), further suggestive of impaired autophagy.

Since NFκB upregulated interleukin expression upon YAP1 targeting/in 3D spheroids and that NFκB suppresses *AR* transcription^46^, we hypothesized that inhibition of pro-survival NFκB signaling may potentiate myCAF deactivation and promote cell death. Supportively, BAY11-7082 mono-treatment modestly increased AR mRNA and protein levels, partially decreased SMA and ITGA11 protein levels and attenuated the upregulation of pro-inflammatory markers induced upon 3D spheroid culture. Importantly, BAY11-7082 further potentiated SB431542/verteporfin/R1881-mediated SMA and ITGA11 downregulation and in the presence of enzalutamide markedly increased cell death (Fig. 6G-H and Supplemental Fig. 9I-J). Collectively, these data suggest that NFκB signaling cooperates with the TGFβ1-YAP1 axis in maintaining the myCAF state, whose deactivation via TGFβ1-YAP1 targeting re-sensitizes them to enzalutamide, most likely via impaired autophagic flux.

## Discussion

Whilst widely considered important therapeutic targets, CAF subpopulations remain poorly characterized at the functional level, primarily due to the paucity of pre-clinical models. The fibroblast biobank generated here contains three distinct fibroblast subtypes: early-activated iCAF associated with a favorable prognosis, an intermediate substate exhibiting molecular hallmarks of both late iCAF and early myCAF, and a subpopulation with a late-activated myCAF profile associated with poor prognostic indicators. Validation in clinical specimens and PDX models underscored the physiological relevance of these distinct subpopulations. The identification of NFκB-TGFβ1-YAP1 signaling as a key driver of stromal AR loss and the aggressive myCAF state provides a mechanistic rationale for adjuvant myCAF targeting to improve castration-resistant PCa response.

The stromal response to tumor presence is not routinely evaluated in pathological tumor staging. Our observation that high-grade PCa manifests as distinct stromal patterns positively (SMC-rich “non-stromogenic”) or negatively (myCAF-rich “stromogenic”) associated with patient survival further supports the inclusion of stromal assessment in diagnostic staging to improve risk stratification^3^. Mechanistically, intact layers of SMC are thought to restrain tumor invasion^47^. Whilst the molecular mechanisms underlying SMC loss in PCa require further investigation, the C3 marker/YAP1 target gene CYR61 is a secreted factor and key inducer of SMC dedifferentiation^48^, implying that myCAF actively contribute to SMC loss. Distinct stromal patterns were also apparent in the two PDX models employed, whereby the myCAF profile of LAPC9 tumors contrasted with the higher expression of early-activation markers in the less aggressive BM18 model. Cancer cell genotype plays a significant role in defining CAF phenotypes e.g., p53 loss enhances stromal fibrosis and myCAF hallmarks^49, 50^. Investigations are underway to determine whether similar genetic aberrations determine stromal patterning in LAPC9/BM18 PDX and clinical PCa. Supportively however, LAPC9 xenografts have a high mutational burden, including *TP53* frameshift mutations, contrasting with the low mutational burden and wild-type *TP53* status of the BM18 model^51^. With regards to the current study, it is noteworthy that p53 modulates extracellular vesicle cargo to dictate specific characteristics of CAF within the TME^52^.

Signaling pathways that delineated biobank culture phenotypes are consistent with the established roles of IL1-JAK-STAT, TNFα-NFκB and TGFβ1 in inducing iCAF and myCAF phenotypes^2, 53^. However, we additionally identify a critical role of YAP1 and NFκB in synergistically acting with TGFβ to maintain the myCAF state, an observation consistent with reports that unidentified stimuli are required in addition to TGFβ1 for complete activation to the myCAF state^22^. Whilst the role of YAP1 in myofibroblast/myCAF conversion is well-established^54^, this is to our knowledge the first time that interaction of these pathways has been demonstrated in the context of myCAF biology.

CAF activation is intimately linked with autophagy, a key process by which cellular components are degraded and recycled to maintain cellular homeostasis^55^. By sequestering substrates to autophagosomes, p62 acts as a selective autophagy receptor and accumulates when autophagy is defective. In this context, YAP transcriptional activity is required for the formation of autolyosomes from autophagosomes^44^. p62 also acts as a signaling hub and activates NFκB promoting the expression of proinflammatory cytokines such as IL6, which promotes TGFβ synthesis^56^. Collectively, findings herein lead us to propose that deactivation of the myofibroblastic state upon TGFβ1-YAP1 inhibition increases demand for autophagic recycling (e.g., of cytoskeletal components) during cellular reprogramming, which however cannot fully ensue due to YAP-impairment. The resulting increase in p62 activates pro-survival NFκB leading to the production of proinflammatory cytokines, including IL6, that could conceivably prime neighboring fibroblasts to undergo myCAF activation. The novel insights provided here into driver mechanisms of myCAF activation and deactivation suggest that therapeutic approaches aiming to reprogram the onco-supportive myCAF state will require combined targeting of the NFκB-TGFβ1-YAP1 axis for persistent myCAF deactivation.

Our observations that androgens promote the proliferation but suppress the migration of early-activated fibroblasts and whose secretome is permissive for PCa cell proliferation imply that under hormone-naïve pre-malignant/low-grade tumor conditions, activated fibroblasts are proliferative but not highly motile. Thus, in the context of fibroblast-led cancer cell migration, such fibroblasts may restrict tumor cell dissemination, providing one potential explanation for the positive association of early-activated fibroblasts with prognostic indicators of favorable outcome. Stromal AR loss in aggressive PCa is poorly understood. We demonstrate that AR loss is associated with the gain of myofibroblastic hallmarks, whereby AR expression was potently reduced by recombinant TGFβ1, and was accompanied by insensitivity to the proliferative effects evoked by modulating AR signaling. Whilst the precise molecular mechanism remains to be determined, it is noteworthy that Smad3 binds to intron 3 of the *AR* gene modulating *AR* expression in PCa cells^57^. However, the findings presented herein collectively implicate the concerted suppressive action of NFκB, TGFβ and YAP1 in contributing to stromal AR loss in PCa.

Our understanding of endogenous stromal AR function in PCa is limited and caution is erred in interpreting findings from ectopic expression models since low levels of endogenous AR in CAF are reportedly transcriptionally-incompetent and elicit an androgenic response profoundly different to the same CAF constitutively overexpressing AR^58^. Despite low AR levels and proliferative-insensitivity to AR modulation, R1881 synergistically downregulated myCAF markers in C3 cultures in combination with SB431542 but not verteporfin, implying specific interaction of AR and TGFβ1 signaling pathways. Intriguingly, enzalutamide in combination with R1881 potentiated myCAF marker downregulation upon TGFβ inhibition alone or in combination with YAP and/or NFκB inhibition. This apparent paradox may arise from AR sequestering and inhibiting the TGFβ signaling intermediate Smad3, similar to that previously reported^57^. We propose that in the context of androgen-mediated AR stabilization, elevated AR levels engage and inhibit Smad3 thereby enhancing SB431542-mediated inhibition of TGFβ signaling. Notably, the AR-Smad3 interaction, which occurs via the AR ligand-binding domain, can proceed in the presence or absence of androgen^57^. Since enzalutamide inhibits ligand binding^59^, it is conceivable that enzalutamide increases availability of the AR ligand-binding domain for Smad3 engagement, thereby potentiating suppressive AR-Smad3 crosstalk.

The aforementioned hypothesis alone however does not explain the marked cell death of SB431542/verteporfin-deactivated myCAF in the presence of R1881 and enzalutamide. This potentially clinically-relevant finding in terms of therapeutic strategies to ablate aggressive myCAF is consistent however with reports that enzalutamide activates AMPK promoting autophagy in PCa cells^60^. In the context of YAP impairment in SB431542/verteporfin-treated myCAF, further promotion of autophagy would be expected to promote detrimental accumulation of autophagosomes resulting in cell death. Further studies are underway to discern the precise molecular mediators underlying enzalutamide-mediated toxicity in deactivated myCAF, however our findings provide a mechanistic rationale for myCAF-targeting adjunct to ADT/ARSI as a potential means to enhance therapeutic efficacy in stromogenic PCa.

ADT constitutes a cornerstone of current systemic treatments for advanced PCa. Ultimately however most patients relapse with lethal castration-resistant disease. We demonstrate that AR^low^/ITGA11^+^/ENG^+^ myCAF, which predominate stromogenic high-grade PCa, have a high propensity to migrate. This is consistent with reports that ITGA11 expression in CAF is required for CAF-induced tumor cell invasion^61^ and that CAF deposit ECM tracks enriched with the C3 marker ITGB5, which serve as guidance cues for cancer cell migration^62^. Together with the proliferative-insensitive nature of C3 to enzalutamide, these findings implicate C3/myCAF in supporting tumor cell dissemination and the development of castration-resistant PCa. Supportively, CAF expressing the C3 markers ENG or SPP1 promoted castration-resistance in PDX and genetically-engineered mouse PCa models, and expanded under castrate conditions^63, 64^. Indeed, we report that C3 myCAF positively correlate with biochemical recurrence in clinical PCa, and were not only prevalent in an aggressive PCa PDX model but were resistant to castration whereupon their myCAF transcriptional profile was further enhanced.

In summary, the data presented here underscore the translational value of our fibroblast biobank for investigating functional aspects of fibroblast heterogeneity in PCa. Moreover, our findings raise important questions with regards to ADT/ARSI that may inadvertently favor survival of the aggressive myCAF substate and imply that adjuvant targeting of the myCAF state at the level of the NFκB-TGFβ-YAP1 axis may improve therapy outcome.

## Methods

### Reagents

Reagents were from Sigma-Aldrich (Vienna, Austria) unless otherwise specified.

### Tissue processing

4 mm^3^ biopsy cores were taken by a Uropathologist (G.S) from freshly excised surgical resections. A small tissue section was removed from the end of each biopsy core, formalin-fixed and paraffin embedded (together with tissue from the surrounding biopsy punch site) for subsequent histopathological evaluation via hematoxylin and eosin (HE) staining and p63/AMACR dual immunohistochemistry (Supplemental Fig. 1). The remaining biopsy core was transferred to cell culture facilities in transport solution (serum-free Dulbecco’s modified eagle media (DMEM, PAN-Biotech GmbH, Aidenbach, Germany) containing 1000 g/L glucose supplemented with 1% penicillin/streptomycin (PAN-Biotech).

### Cell culture

Prostate epithelial cell lines were obtained from American Type Culture Collection (ATCC; Rockville, MD), STR validated and maintained according to the distributor’s instructions. Mycoplasma-free human primary prostatic fibroblasts were established from prostate tissue wedges and maintained as described previously ^65^. Primary fibroblasts were used at passage 8 or lower. All experiments were performed at least three times with primary cells from different donors. In total, this study employed primary fibroblasts isolated from over fifty different donors. All cells were maintained in a humidified atmosphere at 37°C with 5% CO_2_. Cells were maintained in DMEM containing 10% fetal bovine serum (FBS) and supplemented with 1g/L glucose, 1% penicillin/streptomycin and 2% Glutamax (herein termed 10% DMEM). In experiments incorporating R1881 treatment, cells were pre-incubated overnight in DMEM supplemented as above but containing 5% charcoal-treated and thus steroid hormone-depleted FBS (CTFCS) with 10 µM enzalutamide or vehicle equivalent. The following day further treatments were added as indicated in DMEM containing 2.5% CTFBS for the duration stated.

### 3D spheroid formation and matrix stiffness studies

For three-dimensional (3D) fibroblast spheroid generation, 20,000 cells were seeded per well of a Corning® Costar® 96-well round bottom ultra-low attachment plate (Szabo-Scandic HandelsgmbH, Vienna, Austria) in 100 µl fibroblast growth medium and centrifuged at 1200 rpm for 5 min. 18-24 replicates were prepared for each condition. For matrix stiffness studies, 6-well plates were employed containing Cytosoft® hydrogels (Advanced Biomatrix, Sigma-Aldrich) with an elastic modulus of 2 kPa, which is similar to the stiffness of healthy prostate ^66^, and collagen-coated prior to cell seeding.

### Conditioned media generation and application

Fibroblasts were accustomed to reduced serum conditions (2.5% FBS for 12 h) before rinsing in serum-free DMEM (GIBCO^TM^, FisherScientific) and incubated for a further 48 h in serum-free DMEM supplemented with 0.1% BSA, 1% penicillin/streptomycin and 2% GlutaMAX^TM^ (hereon termed SF-DMEM/0.1% BSA) in the presence of R1881, enzalutamide or vehicle equivalent as indicated. The supernatant was collected as conditioned media (CM). Non-CM control media were generated from the same mastermix of SF-DMEM/0.1% BSA but incubated for the same duration without cells. CM were centrifuged at 1000 rpm for 5 min and used immediately or frozen in aliquots for subsequent use. CM were normalized with fresh SF-DMEM/0.1% BSA according to the protein content of lysates from the corresponding cell monolayers as determined via BCA protein assay (Pierce^TM^, Thermo Fisher Scientific, Vienna, Austria). Normalized CM were diluted 2:1 with fresh serum-free media supplemented accordingly for the corresponding tumor cell line and applied to the indicated cells for the duration stated.

### Flow cytometry and fluorescence-activated cell sorting (FACS)

Cells were detached from vessels using TrypLE (Invitrogen), stopped by dilution with PBS and after centrifugation, the pellet resuspended in 200 µl PBS containing 0.5% BSA and 2 mM EDTA (hereon flow cytometry (FC) buffer). 3 x 10^5^ cells were transferred to an Eppendorf tube and resuspended in FC buffer. 3 µl of Intratect 50 g/l infusion solution (Biotest AG, Dreieich, Germany) was added and samples incubate on ice for 5 min. Antibodies and BD Horizon™ Brilliant Stain Buffer Plus (BD Bioscience) were added as indicated (Supplemental Table 13) and samples incubated for 30 min at 4°C protected from light. FC buffer (for FACS samples) or BD Pharm Lyse™ Lysing Buffer (for FC; BD Bioscience) was added to a final volume of 2 ml and cells pellet by centrifugation after incubation for 10 min at RT. Cell suspensions were washed twice with 2 ml FC buffer and cells resuspended in 100 µl FC buffer and maintained at 4°C until measurement. Approximately 5 min prior to analysis, 4 µl BD Pharmingen™ 7-AAD viability dye (BD Bioscience) was added to each sample. Flow cytometry was performed on a FACSymphony™ A5 cell analyzer (BD Biosciences) equipped with 355 nm, 407 nm, 561 nm, and 639 nm laser lines and running BD FACSDiva (v9.1) software. All antibodies were titrated in previous experiments to determine optimal staining concentration in 100µl cell suspensions. Unstained and fluorescence-minus-one (FMO) samples were conducted for gate setting during data analysis.

For FACS, cultures were processed as above with reagent volumes adjusted for a final volume of 500 µl. After staining and washing, cells were resuspended in sorting buffer (HBSS containing 0.5% BSA, 2 mM EDTA and 25 mM HEPES). After the addition of 7AAD viability dye, samples were strained though a 100 µm mesh filter and viable cells sorted on a FACS Aria I (BD Biosciences) for cluster 1 and clusters 6 and 10 per the gating strategy outlined in Supplemental Fig. 5D whereby in one experiment (lower panel, Supplemental Fig. 5I) cells corresponding to clusters 6 and 10 were pooled into a single sample. Cells were sorted directly into Rlyse buffer for RNA isolation.

### Flow cytometry data analysis

FCS files were analyzed using FlowJo™ (BD Life Sciences, v10.10.0). Cells were gated based on size and viability stain (FSC-A/7-AAD) to exclude debris/dead cells and doublets removed (FSC-A/FSC-H) generating a single viable cell population for analysis. FC counts from all samples (n=23) were concatenated into a single file, which retained the original sample labelling. Using the t-distributed stochastic neighbour embedding (tSNE) option in FlowJo™ dimensional reduction was performed based on seven stromal markers (MCAM, CD140a, CD140b, CD90, PDPN, FAP, CD105). The learning configuration was set to auto (opt-SNE), with 1000 iterations, a perplexity of 30, and a learning rate set to 1/12 of the count of the concatenated file. The Exact (vantage point tree) KNN algorithm and the FFT Interpolation (FIt-SNE) gradient algorithm were used. Putative subclusters were calculated with XShift based on the seven stromal markers, using 500 as the number of nearest neighbours (K), the Euclidean distance metric, a subsampling limit of 10^5, and an auto Run ID. Identified clusters were examined using ClusterExplorer and manual gating. Unstained and fluorescence-minus-one (FMO) samples were used to define gating thresholds for each stromal marker, forming the basis of the gating strategy. The developed gating strategy was then applied to additional samples (n=13) to analyze gated clusters composition.

### RNA isolation, bulk transcriptomics, cDNA synthesis and quantitative real time PCR (qRT-PCR)

Total RNA isolation, cDNA synthesis and qRT-PCR using Taqman® gene expression assays were performed as described^65^. RNA isolation of FACS-sorted cells was performed using the RNeasy Micro Plus kit (Qiagen GmbH, Hamburg, Germany) according to the manufacturer’s instructions. TaqMan® gene expression assays (Thermo Fisher Scientific) are listed in Supplemental Table S11. One microgram total RNA was hybridized to whole genome BeadChip® Sentrix arrays (HumanHT-12v4) at the German Cancer Genome Research Centre, Heidelberg, Germany). Raw data were processed as described^67^ and differential gene expression analyses performed on quantile normalized data using limma^68^. Genes displaying *P.*adj<0.05 and log2FC>0.3 were considered upregulated.

### Bioinformatics

Bioinformatic analysis was performed in R (v4.2.1) using both base R^69^ and Bioconductor^70^ packages. Fibroblast clusters were identified using principal component analysis (PCA) and hierarchical clustering using log_e_(1+ quantile normalized count) with the Euclidian distance used to calculate the distance matrix according to Ward’s linkage^71^. Comparison to publicly available fibroblast/CAF signatures was performed by calculating combined z-scores ^72^ using the tidy framework hacksig^73^.

Gene expression or combined z-scores were visualized using EnhancedVolcano^74^ and/or pheatmap^75^. Pathway analysis was performed using clusterProfiler^76^, msigdbr^77^ and Pathway RespOnsive GENes (PROGENy) for activity inference^78^. YAP1 target genes were defined as reported^79^. R clustering of FC data was performed in R (v4.2.1) using RStudio (v2022.7.1.554) incorporating base R functionalities^69^ and additional packages. Data manipulation was carried out with the tidyverse package (v2.0.0) ^80^. Heatmaps were generated using the pheatmap package (v1.0.12) ^75^, with color gradients applied via the RColorBrewer package (v1.1-3) ^81^. The percentages of the gated clusters in each sample were visualized as a heatmap and scaled per sample to highlight the most prominent clusters in each sample, facilitating the grouping of samples into broader categories. The percentages of the original clusters positive for each of the seven stromal markers or the C1, C2 or C3 subpopulations were visualized as a heatmap without scaling.

### Analysis of TCGA and SU2C data

RNA sequencing and clinical data of TCGA prostate adenocarcinoma (PRAD) ^82^ samples were downloaded from UCSC Xena (https://xenabrowser.net/datapages/). Combined z-scores were calculated for bulk transcriptomic-derived gene signatures as described above and compared between clinical parameters. The R package ggsignif^83^ was used to perform t-tests. For visualization as heatmaps, only samples with the sample type “primary tumor” were selected. Disease-free survival (DFS) analyses were performed using GEPIA2^84^ using the group cut-offs indicated in the corresponding figure legend. Bulk transcriptomic-derived signatures were generated from the top 5 marker genes for each fibroblast cluster ranked according to *P.*adj value with log2 fold change (log2FC) >0.5. Gene signatures are provided in the Source Data file. The SU2C metastatic prostate adenocarcinoma dataset^85^ was analyzed using cBioportal^86^. Samples were divided based on expression of the top 15 CAF-subtype specific markers using OQL. A sample was considered “high” for one CAF subtype if 3 or more CAF subtype-specific markers were > z-score of 1 (number of samples/patients: C1 = 93/93, C2 = 103/101, C3 = 93/91). Samples (90) that overlap in the selected groups were excluded from sample-level analysis which resulted in 34, 29 and 24 samples for C1, C2 and C3, respectively. Clinical attributes were compared between those groups. For analysis of SMC gene expression on survival in high-grade PCa, the TCGA-PRAD cohort (n=500) was initially stratified into high-(≥T2c) and low-grade (≤T2b) tumors based on pathological T stage. The data were subsequently filtered according to the following criteria: 2 patients were excluded due to prior treatment, and 29 patients with low-grade tumors were excluded from the survival analysis. The top 20 upregulated genes from prostate SMC gene signatures were curated from relevant literature^16, 87^ and are provided in the source data file. These gene lists were condensed into gene signature expression values using the GSVA R package (v1.50.5). The filtered TCGA PRAD cohort (n = 469) was then stratified into “high” and “low” SMC expression groups, with the threshold determined by the optimal cutpoint calculated using the surv_cutpoint() function from the survminer R package (v0.4.9). Survival analysis was conducted using the survival R package (v3.7-0), and Kaplan-Meier curves were generated with the ggsurvplot() function from survminer (v0.4.9).

### Cell size analysis

Subconfluent fibroblast cultures were trypsinized and cell size determined on a CASY cell counter and analyzer (Schärfe-System GmbH, Reutlingen, Germany) equipped with a 150 µm capillary using the following settings: evaluation cursors 11.18 – 50.00 µm, normalization cursors 6.88 – 50.00 µm, smoothing 0 and automatic aggregation correction.

### SDS-PAGE and Western blotting

Preparation of cell lysates, normalization via BCA protein assay (Pierce, Thermo Scientific), SDS-PAGE and Western blotting were performed as described^65^ using antibodies as indicated in Supplemental Table S12.

### Proliferation assays

Cells seeded in quadruplicate in black 96-well plates in phenol red-free media were treated for 96h as indicated. WST-1 assays (Sigma Aldrich) were performed according to the manufacturer’s instructions before absorbance measurement at 490 nm on a BioTek Cytation 5 plate reader (Agilent Technologies Österreich GmbH, Vienna, Austria) running BioTek Gen5 software (version 3.09, Agilent Technologies Österreich GmbH). Background intensity from cell-free wells was subtracted. Thereafter media were aspirated and cells lyzed by freezing at −80°C. Upon thawing, 200 μl CyQuant lysis reagent (Invitrogen, Vienna, Austria) was added containing SYBR® Green I (diluted 1:1000). Following incubation for 30 min at 37°C, fluorescence was measured on a BioTek Cytation 5 plate reader (Agilent Technologies Österreich GmbH) and background intensity from cell-free wells subtracted. A replica seeded plate harvested immediately upon cell attachment (<4h) was used to normalize potential differences in seeding densities between different fibroblast cultures. Real time proliferation assays were conducted as above but analyzed on an Incucyte ® S3 Live-cell analysis instrument (Sartorius Lab Instruments GmbH and Co. KG, Gladbach, Germany). Images were acquired every 12 h and the phase object confluence determined.

### Migration assays

Migration assays using 24-well Fluoroblok transwell plates with an 8 μm pore size (BD Bioscience, Vienna, Austria) were performed as described^65^ with the exception that fluorescence was measured on a BioTek Cytation 5 plate reader (Agilent Technologies Österreich GmbH) in bottom read mode.

### Patient-derived xenografts (PDX)

The bone metastatic PCa PDX models have been previously described^88, 89^. BM18 and LAPC9 PDX FFPE tissue blocks and bulk RNA-sequencing data (European Genome-Phenome Archive, Accession number EGAS00001004770) used herein derived from our previous study^35^.

### Immunocytochemistry and immunohistochemistry

For immunohistochemistry, 2 µm FFPE tissue sections from consenting treatment-naive patients were stained on a Ventana Benchmark Ultra automated staining device (Roche Diagnostics GmbH, Vienna, Austria) with the antibodies denoted in Supplemental Table S13. For multiplex immunofluorescent staining, tissue sections were deparaffinized and rehydrated in a graded alcohol series. Antigen retrieval was performed via indirect boiling in Dako Target Retrieval Solution pH 9 (Agilent) for 10 min.

After cooling, sections were blocked in 3% BSA in Tris-buffered saline (TBS) before sequential incubation for 1h at RT or overnight at 4°C with primary antibodies diluted in 0.5% BSA in TBS as indicated (Supplemental Table 13). Washed sections were incubated with fluorescently-conjugated secondary antibodies (Supplemental Table 13) for 1h at RT. The cycle was repeated until all antibodies had been stained. Nuclei were counterstained with 2.5 µg/ml Hoechst 33342 (Invitrogen, Thermofisher) before mounting in VECTASHIELD® mounting medium for fluorescence (Vector Laboratories, Inc., Burlington, CA). Images were acquired on a Zeiss Axio Imager Z2 microscope (Zeiss, Vienna, Austria) equipped with a Pixelink PL-B622-CU camera for brightfield imaging and a monochrome pco.edge 4.2LT camera for fluorescence imaging. TissueFAXS® software (version 7.137, TissueGnostics® GmbH, Vienna, Austria) was used to acquire images via a 20x air objective using constant image acquisition settings. Single channel monochrome images were merged and pseudo-colored for visualization as indicated in the corresponding figure legend. Quantification of immunofluorescent images were performed in ImageJ 1.54j (Rasband, NIH, USA). Masks were created to calculate the DAPI^+^ area (all nuclei), DAPI and AR area (AR^+^ nuclei) and applied to masks created in the stromal marker-specific channel to select for the DAPI^+^AR^+^ area for and each stromal marker. Data are shown as a percentage of all AR^+^ nuclei positive for each stromal cell marker relative to the total area positive for DAPI and the corresponding stromal cell marker.

### Statistics and reproducibility

Unless otherwise specified cell line data are shown as mean ± SEM. *n* represents the number of biological replicates as stated in the corresponding figure legends. All experiments were repeated independently at least three times using primary fibroblasts isolated from different donors. Statistical analyses were performed using the R statistical environment (v4.2.1) ^69^ or GraphPad Prism (v9.5.0 GraphPad Software, LLC). Methods used for each statistical analysis are specified in the corresponding figure legend or relevant section of the Materials and Methods. Unless otherwise specified, statistical significance is denoted: NS, not significant; *, *P*<0.05; **, *P*<0.01; ***, *P*<0.001.

## Declarations

### Ethics approval and consent to participate

Prostate tissue samples were collected from consenting patients undergoing radical prostatectomy at the University Hospital of Innsbruck in accordance with the prior approval from the ethics committee of the Medical University of Innsbruck (1129/2022). Use of archived patient-derived FFPE samples was approved by the ethics committee of the Innsbruck Medical University (1072/2018, UN4837). All patients gave written informed consent.

### Consent for publication

All authors have read and consent to this publication.

### Availability of data, materials and code

Human bulk transcriptomic explant culture dataset is available in ArrayExpress under accession number E-MTAB-13167. Mouse RNA-sequencing reads for the PDX models are available in the European Genome–Phenome Archive database under accession number EGAS00001004770. Source data are provided in the supplementary file. No new code was developed for this manuscript.

## Competing interests

The authors have no competing interests.

## Funding

This work was funded by the Austrian Science Fund (FWF, P31122 and I4565) and the Swiss National Science Fund (SNSF, 189369 and 189149) and co-funded by the Tiroler Wissenschaftsfond.

## Authors’ contributions

Conception or design: S.K., M.K. and N.S.

Data acquisition, analysis or interpretation: E.B., M.G., E.D., G.F., G.S., L. Nommensen, L. Neumann, S.K., F.B., M.K. and N.S.

Writing: E.B., M.K. and N.S.

Supervision and funding acquisition: M.K. and N.S.

## Acknowledgements

The authors gratefully acknowledge Dr. Gabriele Dobler for assistance with cell culture, Eberhard Steiner for retrieval of clinical data, Matthias Vianello for assistance with western blotting and Géza Tamás Szabó for bioinformatic analyses. The results shown here are in whole or part based upon data generated by the TCGA Research Network (https://www.cancer.gov/tcga). Schematic illustrations were created using BioRender.com.

## Abbreviations

2D: two-dimensional
3D: three-dimensional
ADT: androgen deprivation therapy
apCAF: antigen-presenting cancer-associated fibroblast
AR: androgen receptor
ARSI: androgen receptor signaling inhibitor
BSA: bovine serum albumin
CAF: cancer-associated fibroblasts
CES1: carboxylesterase I
CM: conditioned media
CTFBS: charcoal treated fetal bovine serum
DFS: disease-free survival
DMEM: Dulbecco’s modified eagle media
ECM: extracellular matrix
ENG: endoglin
FAP: fibroblast activation protein
FBS: fetal bovine serum
FFPE: formalin-fixed paraffin-embedded
GO: gene ontology
h: hours
HBSS: Hank’s balanced salt solution
HE: hematoxylin and eosin
HGPIN: high-grade prostatic intraepithelial neoplasia
iCAF: inflammatory cancer-associated fibroblast
min: minutes
myCAF: myofibroblastic cancer-associated fibroblast
nonCM: non-conditioned media
PAAD: pancreatic ductal adenocarcinoma
PAGE4: prostate-associated gene 4
PCA: principal component analysis
PCa: prostate cancer
PDX: patient-derived xenograft
PFS: progression-free survival
PIN: prostatic intraepithelial neoplasia
PRAD: prostate adenocarcinoma
PSA: prostate-specific antigen
qRT-PCR: quantitative real time PCR
rpm: revolutions per minute
RT: room temperature
scRNA-seq: single cell RNA sequencing
SEM: standard error of the mean
SF-DMEM: serum-free Dulbecco’s modified eagle medium
SMA: alpha smooth muscle actin
SMC: smooth muscle cells
STR: short tandem repeat
TCP: standard tissue culture plasticware
TGFβ: transforming growth factor beta
TME: tumor microenvironment
YAP1: yes-associated protein 1

## Notes

### Competing Interest Statement

The authors have declared no competing interest.

## References

1. Pederzoli F, Raffo M, Pakula H, Ravera F, Nuzzo PV, Loda M. “Stromal cells in prostate cancer pathobiology: friends or foes?”. Br J Cancer 128, 930–939 (2023).

2. Cheng PSW, Zaccaria M, Biffi G. Functional heterogeneity of fibroblasts in primary tumors and metastases. Trends Cancer, (2024).

3. Ruder S, et al. Development and validation of a quantitative reactive stroma biomarker (qRS) for prostate cancer prognosis. Hum Pathol 122, 84–91 (2022).

4. Rebello RJ, et al. Prostate cancer. Nat Rev Dis Primers 7, 9 (2021).

5. Leach DA, Fernandes RC, Bevan CL. Cellular specificity of androgen receptor, coregulators, and pioneer factors in prostate cancer. Endocr Oncol 2, R112–R131 (2022).

6. Liu Y, et al. Stromal AR inhibits prostate tumor progression by restraining secretory luminal epithelial cells. Cell Rep 39, 110848 (2022).

7. Cioni B, et al. Loss of androgen receptor signaling in prostate cancer-associated fibroblasts (CAFs) promotes CCL2- and CXCL8-mediated cancer cell migration. Mol Oncol 12, 1308–1323 (2018).

8. Welsh M, et al. Smooth muscle cell-specific knockout of androgen receptor: a new model for prostatic disease. Endocrinology 152, 3541–3551 (2011).

9. Lavie D, Ben-Shmuel A, Erez N, Scherz-Shouval R. Cancer-associated fibroblasts in the single-cell era. Nat Cancer 3, 793–807 (2022).

10. Croizer H, et al. Deciphering the spatial landscape and plasticity of immunosuppressive fibroblasts in breast cancer. Nat Commun 15, 2806 (2024).

11. Cords L, et al. Cancer-associated fibroblast classification in single-cell and spatial proteomics data. Nat Commun 14, 4294 (2023).

12. Buechler MB, et al. Cross-tissue organization of the fibroblast lineage. Nature 593, 575–579 (2021).

13. Nicolas AM, et al. Inflammatory fibroblasts mediate resistance to neoadjuvant therapy in rectal cancer. Cancer Cell 40, 168–184 e113 (2022).

14. Hanley CJ, et al. Single-cell analysis reveals prognostic fibroblast subpopulations linked to molecular and immunological subtypes of lung cancer. Nat Commun 14, 387 (2023).

15. Kieffer Y, et al. Single-Cell Analysis Reveals Fibroblast Clusters Linked to Immunotherapy Resistance in Cancer. Cancer Discov 10, 1330–1351 (2020).

16. Chen Y, et al. Single-cell transcriptomics reveals cell type diversity of human prostate. J Genet Genomics 49, 1002–1015 (2022).

17. Mathieson L, Koppensteiner L, Dorward DA, O’Connor RA, Akram AR. Cancer-associated fibroblasts expressing fibroblast activation protein and podoplanin in non-small cell lung cancer predict poor clinical outcome. Br J Cancer 130, 1758–1769 (2024).

18. Neuzillet C, et al. Periostin- and podoplanin-positive cancer-associated fibroblast subtypes cooperate to shape the inflamed tumor microenvironment in aggressive pancreatic adenocarcinoma. J Pathol 258, 408–425 (2022).

19. Jenkins BH, Buckingham JF, Hanley CJ, Thomas GJ. Targeting cancer-associated fibroblasts: Challenges, opportunities and future directions. Pharmacol Ther 240, 108231 (2022).

20. Costa A, et al. Fibroblast Heterogeneity and Immunosuppressive Environment in Human Breast Cancer. Cancer Cell 33, 463–479 e410 (2018).

21. Elyada E, et al. Cross-Species Single-Cell Analysis of Pancreatic Ductal Adenocarcinoma Reveals Antigen-Presenting Cancer-Associated Fibroblasts. Cancer Discov 9, 1102–1123 (2019).

22. Jenkins BH, et al. Single cell and spatial analysis of immune-hot and immune-cold tumours identifies fibroblast subtypes associated with distinct immunological niches and positive immunotherapy response. Mol Cancer 24, 3 (2025).

23. Koncina E, et al. IL1R1(+) cancer-associated fibroblasts drive tumor development and immunosuppression in colorectal cancer. Nat Commun 14, 4251 (2023).

24. Gao Y, et al. Cross-tissue human fibroblast atlas reveals myofibroblast subtypes with distinct roles in immune modulation. Cancer Cell 42, 1764–1783 e1710 (2024).

25. Schoonderwoerd MJA, Goumans MTH, Hawinkels L. Endoglin: Beyond the Endothelium. Biomolecules 10, (2020).

26. Erdogan B, et al. Cancer-associated fibroblasts promote directional cancer cell migration by aligning fibronectin. J Cell Biol 216, 3799–3816 (2017).

27. Li G, et al. Interfering with lipid metabolism through targeting CES1 sensitizes hepatocellular carcinoma for chemotherapy. JCI Insight 8, (2023).

28. Vandooren J, Itoh Y. Alpha-2-Macroglobulin in Inflammation, Immunity and Infections. Front Immunol 12, 803244 (2021).

29. Biffi G, et al. IL1-Induced JAK/STAT Signaling Is Antagonized by TGFbeta to Shape CAF Heterogeneity in Pancreatic Ductal Adenocarcinoma. Cancer Discov 9, 282–301 (2019).

30. Ahuja S, Sureka N, Zaheer S. Unraveling the intricacies of cancer-associated fibroblasts: a comprehensive review on metabolic reprogramming and tumor microenvironment crosstalk. APMIS 132, 906–927 (2024).

31. Leach DA, Buchanan G. Stromal Androgen Receptor in Prostate Cancer Development and Progression. Cancers (Basel) 9, (2017).

32. Ohlund D, et al. Distinct populations of inflammatory fibroblasts and myofibroblasts in pancreatic cancer. J Exp Med 214, 579–596 (2017).

33. Lumahan LEV, Arif M, Whitener AE, Yi P. Regulating Androgen Receptor Function in Prostate Cancer: Exploring the Diversity of Post-Translational Modifications. Cells 13, (2024).

34. Wu F, et al. Signaling pathways in cancer-associated fibroblasts and targeted therapy for cancer. Signal Transduct Target Ther 6, 218 (2021).

35. Karkampouna S, et al. Stroma Transcriptomic and Proteomic Profile of Prostate Cancer Metastasis Xenograft Models Reveals Prognostic Value of Stroma Signatures. Cancers (Basel) 12, (2020).

36. Maykel J, Liu JH, Li H, Shultz LD, Greiner DL, Houghton J. NOD-scidIl2rg (tm1Wjl) and NOD-Rag1 (null) Il2rg (tm1Wjl): a model for stromal cell-tumor cell interaction for human colon cancer. Dig Dis Sci 59, 1169–1179 (2014).

37. Zhu ML, Kyprianou N. Androgen receptor and growth factor signaling cross-talk in prostate cancer cells. Endocr Relat Cancer 15, 841–849 (2008).

38. Shen T, et al. YAP1 plays a key role of the conversion of normal fibroblasts into cancer-associated fibroblasts that contribute to prostate cancer progression. J Exp Clin Cancer Res 39, 36 (2020).

39. Rosenbluh J, et al. beta-Catenin-driven cancers require a YAP1 transcriptional complex for survival and tumorigenesis. Cell 151, 1457–1473 (2012).

40. Guo Y, Luo J, Zou H, Liu C, Deng L, Li P. Context-dependent transcriptional regulations of YAP/TAZ in cancer. Cancer Lett 527, 164–173 (2022).

41. Ghomlaghi M, et al. Integrative modeling and analysis of signaling crosstalk reveal molecular switches coordinating Yes-associated protein transcriptional activities. iScience 27, 109031 (2024).

42. Levy D, Adamovich Y, Reuven N, Shaul Y. Yap1 phosphorylation by c-Abl is a critical step in selective activation of proapoptotic genes in response to DNA damage. Mol Cell 29, 350–361 (2008).

43. Wei J, et al. The role of matrix stiffness in cancer stromal cell fate and targeting therapeutic strategies. Acta Biomater 150, 34–47 (2022).

44. Sari D, Gozuacik D, Akkoc Y. Role of autophagy in cancer-associated fibroblast activation, signaling and metabolic reprograming. Front Cell Dev Biol 11, 1274682 (2023).

45. Moscat J, Karin M, Diaz-Meco MT. p62 in Cancer: Signaling Adaptor Beyond Autophagy. Cell 167, 606–609 (2016).

46. Tahsin S, et al. AR loss in prostate cancer stroma mediated by NF-kappaB and p38-MAPK signaling disrupts stromal morphogen production. Oncogene 43, 2092–2103 (2024).

47. Liu W, Wang M, Wang M, Liu M. Single-cell and bulk RNA sequencing reveal cancer-associated fibroblast heterogeneity and a prognostic signature in prostate cancer. Medicine (Baltimore) 102, e34611 (2023).

48. Harryman WL, Marr KD, Hernandez-Cortes D, Nagle RB, Garcia JGN, Cress AE. Cohesive cancer invasion of the biophysical barrier of smooth muscle. Cancer Metastasis Rev 40, 205–219 (2021).

49. Chhabra Y, Weeraratna AT. Fibroblasts in cancer: Unity in heterogeneity. Cell 186, 1580–1609 (2023).

50. Wormann SM, et al. Loss of P53 Function Activates JAK2-STAT3 Signaling to Promote Pancreatic Tumor Growth, Stroma Modification, and Gemcitabine Resistance in Mice and Is Associated With Patient Survival. Gastroenterology 151, 180–193 e112 (2016).

51. Karkampouna S, et al. Patient-derived xenografts and organoids model therapy response in prostate cancer. Nat Commun 12, 1117 (2021).

52. Naito Y, Yoshioka Y, Ochiya T. Intercellular crosstalk between cancer cells and cancer-associated fibroblasts via extracellular vesicles. Cancer Cell Int 22, 367 (2022).

53. Hanley C, et al. Spatially discrete signalling niches regulate fibroblast heterogeneity in human lung cancer. bioRxiv, 2020.2006.2008.134270 (2020).

54. Baroja I, Kyriakidis NC, Halder G, Moya IM. Expected and unexpected effects after systemic inhibition of Hippo transcriptional output in cancer. Nat Commun 15, 2700 (2024).

55. Kumar AV, Mills J, Lapierre LR. Selective Autophagy Receptor p62/SQSTM1, a Pivotal Player in Stress and Aging. Front Cell Dev Biol 10, 793328 (2022).

56. Valencia T, et al. Metabolic reprogramming of stromal fibroblasts through p62-mTORC1 signaling promotes inflammation and tumorigenesis. Cancer Cell 26, 121–135 (2014).

57. Chipuk JE, et al. The androgen receptor represses transforming growth factor-beta signaling through interaction with Smad3. J Biol Chem 277, 1240–1248 (2002).

58. Di Donato M, et al. The androgen receptor/filamin A complex as a target in prostate cancer microenvironment. Cell Death Dis 12, 127 (2021).

59. Ito Y, Sadar MD. Enzalutamide and blocking androgen receptor in advanced prostate cancer: lessons learnt from the history of drug development of antiandrogens. Res Rep Urol 10, 23–32 (2018).

60. Elshazly AM, Gewirtz DA. Making the Case for Autophagy Inhibition as a Therapeutic Strategy in Combination with Androgen-Targeted Therapies in Prostate Cancer. Cancers (Basel) 15, (2023).

61. Primac I, et al. Stromal integrin alpha11 regulates PDGFR-beta signaling and promotes breast cancer progression. J Clin Invest 129, 4609–4628 (2019).

62. Baschieri F, et al. Fibroblasts generate topographical cues that steer cancer cell migration. Sci Adv 9, eade2120 (2023).

63. Wang H, et al. Antiandrogen treatment induces stromal cell reprogramming to promote castration resistance in prostate cancer. Cancer Cell 41, 1345–1362 e1349 (2023).

64. Kato M, et al. Heterogeneous cancer-associated fibroblast population potentiates neuroendocrine differentiation and castrate resistance in a CD105-dependent manner. Oncogene 38, 716–730 (2019).

65. Sampson N, et al. Inhibition of Nox4-dependent ROS signaling attenuates prostate fibroblast activation and abrogates stromal-mediated protumorigenic interactions. Int J Cancer 143, 383–395 (2018).

66. Hoyt K, et al. Tissue elasticity properties as biomarkers for prostate cancer. Cancer Biomark 4, 213–225 (2008).

67. Rattay K, Derbinski J, Hofmann TG, Kyewski B. Genome-wide gene expression profiling of homeodomain-interacting protein kinase 2 deficient medullary thymic epithelial cells. Genom Data 6, 48–50 (2015).

68. Ritchie ME, et al. limma powers differential expression analyses for RNA-sequencing and microarray studies. Nucleic Acids Res 43, e47 (2015).

69. R Core Team. R: A language and environment for statistical computing.). R Foundation for Statistical Computing (2022).

70. Huber W, et al. Orchestrating high-throughput genomic analysis with Bioconductor. Nat Methods 12, 115–121 (2015).

71. Ward JH. Hierarchical Grouping to Optimize an Objective Function. J Am Stat Assoc 58, 236-& (1963).

72. Lee E, Chuang HY, Kim JW, Ideker T, Lee D. Inferring pathway activity toward precise disease classification. PLoS Comput Biol 4, e1000217 (2008).

73. Carenzo A, Pistore F, Serafini MS, Lenoci D, Licata AG, De Cecco L. hacksig: a unified and tidy R framework to easily compute gene expression signature scores. Bioinformatics 38, 2940–2942 (2022).

74. Blighe K, Rana S, Lewis M. EnhancedVolcano: Publication-ready volcano plots with enhanced colouring and labeling.). R package version 1.16.0 edn (2022).

75. Kolde R. pheatmap: Pretty Heatmaps.). R package version 1.0.12 edn (2019).

76. Wu T, et al. clusterProfiler 4.0: A universal enrichment tool for interpreting omics data. Innovation (Camb) 2, 100141 (2021).

77. Dolgalev I. msigdbr: MSigDB Gene Sets for Multiple Organisms in a Tidy Data Format.). R package version 7.5.1 edn (2022).

78. Schubert M, et al. Perturbation-response genes reveal signaling footprints in cancer gene expression. Nat Commun 9, 20 (2018).

79. Wang Y, et al. Comprehensive Molecular Characterization of the Hippo Signaling Pathway in Cancer. Cell Rep 25, 1304–1317 e1305 (2018).

80. Wickham H, et al. Welcome to the Tidyverse. Journal of Open Source Software 4, 1686–1693 (2019).

81. Neuwirth E. _RColorBrewer: ColorBrewer Palettes_.) (2022).

82. Cancer Genome Atlas Research N. The Molecular Taxonomy of Primary Prostate Cancer. Cell 163, 1011–1025 (2015).

83. Ahlmann-Eltze C. GGSIGNIF: R package for displaying significance brackets for ‘GGPLOT2’.). 2 edn (2021).

84. Tang Z, Kang B, Li C, Chen T, Zhang Z. GEPIA2: an enhanced web server for large-scale expression profiling and interactive analysis. Nucleic Acids Res 47, W556–W560 (2019).

85. Abida W, et al. Genomic correlates of clinical outcome in advanced prostate cancer. Proc Natl Acad Sci U S A 116, 11428–11436 (2019).

86. Cerami E, et al. The cBio cancer genomics portal: an open platform for exploring multidimensional cancer genomics data. Cancer Discov 2, 401–404 (2012).

87. Joseph DB, et al. Single-cell analysis of mouse and human prostate reveals novel fibroblasts with specialized distribution and microenvironment interactions. J Pathol 255, 141–154 (2021).

88. McCulloch DR, Opeskin K, Thompson EW, Williams ED. BM18: A novel androgen-dependent human prostate cancer xenograft model derived from a bone metastasis. Prostate 65, 35-43 (2005).

89. Craft N, et al. Evidence for clonal outgrowth of androgen-independent prostate cancer cells from androgen-dependent tumors through a two-step process. Cancer Res 59, 5030–5036 (1999).

90. Heidegger I, et al. Comprehensive characterization of the prostate tumor microenvironment identifies CXCR4/CXCL12 crosstalk as a novel antiangiogenic therapeutic target in prostate cancer. Mol Cancer 21, 132 (2022).

