## Supplemental Figures and Legends for "Unraveling the YAP1-TGFβ1 axis: a key driver of androgen receptor loss in prostate cancer-associated fibroblasts"

**Supplemental figure legends**

**Supplemental Figure 1 Isolation and characterization of *ex vivo* culture of primary prostate fibroblasts**

A) RP tissue wedges (i) containing regions of suspected malignancy and benign-adjacent tissue are sampled (ii) using 4 $mm^3$ biopsy punchers. The top part of each biopsy core is removed (iii) for FFPE processing (iv) and subsequent histopathological evaluation. Remaining tissue is quartered (v) and similarly processed. B) Histopathological validation of the top section of biopsy cores from (Aiii-iv) via HE staining and p63 (brown) and AMACR (red) dual-immunohistochemistry. C) The remaining biopsy core is cut into small pieces and transferred to culture medium that supports fibroblast but not endothelial cell growth. Outgrowing fibroblasts are selectively enriched away via trypsinization from any epithelial cells, which require additional collagenase treatment for detachment. D) Representative images of three primary fibroblast explant cultures isolated from different patients stained for mesenchymal markers vimentin (green) and CD90 (white) and the epithelial marker pan-cytokeratin (red). Nuclei were counterstained using Hoechst (blue). 22Rv1 PCa cells served as positive control for pan-cytokeratin (*far right, upper panel).* Negative control of a parallel stained fibroblast culture incubated without primary antibodies (*far right, lower panel).*

**Supplemental Figure 2 *Ex vivo* culture of primary prostate fibroblasts – pertaining to Supplemental Fig. 1**

Single channel monochromatic images (from Supplemental Fig. 1D) of immunofluorescent validation of three primary fibroblast explant cultures isolated from different patients stained for mesenchymal markers vimentin (green) and CD90 (white) and the epithelial marker pan-cytokeratin (pan CK, red). Nuclei were counterstained using Hoechst (blue). 22Rv1 PCa cells served as a positive control (pos. ctrl) for pan-cytokeratin. Negative control (neg. ctrl) of a parallel stained fibroblast culture incubated without primary antibodies (*far right, lower panel).*

**Supplemental Figure 3 Transcriptomic analysis of primary prostate fibroblast cultures identifies two distinct CAF populations - pertaining to Figure 1**

A) Heatmap depicting sample level expression of significantly upregulated genes (*P*.adj<0.05, log2FC>0.3) for each fibroblast cluster in the transcriptomic dataset (related to Fig. 1C). B) clusterProfiler analysis of *left:* GO Biological Process for significantly upregulated genes of each fibroblast cluster, including two cycling CAF explant cultures (cycling), *centre and right*: the top 20 scoring Hallmark and REACTOME pathways for significantly upregulated genes of each fibroblast cluster (related to Fig. 1E-G). C-D) Expression of the indicated genes across each fibroblast cluster. Values denote C) average log2 signal intensity from quartile normalized bulk transcriptomic data and D) mean $2^{-\Delta\Delta Ct}$ via qRT-PCR relative to the housekeeping gene *TBP*. Error bars denote S.E.M. Significance was calculated via a Kruskal-Wallis test with multi-comparison correction using the two-stage linear step-up method of Benjamini, Krieger and Yekutieli. E) Comparison of expression profiles for each fibroblast cluster with published scRNA-seq datasets using combined z-scores for each gene signature and represented as mean per fibroblast cluster (related to Fig. 1I-J). F) Expression of signature genes (combined z-score) for each primary fibroblast cluster in the TCGA prostate adenocarcinoma (PRAD) cohort (related to Fig. 1K-N). Boxplots were used to compare signature gene expression levels across Gleason scores, T and N stages and biochemical recurrence. The R package ggsignif [74] was used to perform t-tests. G) Expression levels of substate-delineating markers used to perform survival analyses (Fig. 1M) in different cell types of a PCa scRNA-seq dataset[78]. Source data for panels A-C, E-F are provided in the Source Data file.

**Supplemental Figure 4 Primary explant cultures depict distinct stable homogeneous fibroblast phenotypes**

A) Sample level heatmap (clustered according to ward.D) for qRT-PCR-derived $2^{-\Delta\Delta CT}$ values of the indicated genes in an independent cohort of 29 primary human prostate explant cultures (qRT-PCR cohort) and fresh cultures from 3 donors used in the original bulk transcriptomic cohort. Sample numbers are indicated beneath the heatmap. Font color denotes the subpopulation annotated using the different methodologies (C1, blue; C2, yellow; C3, red; n.d., not determined). B) Expression level of the indicated genes in patient-matched C1 and C2 cultures maintained longitudinally for the indicated number of passages. Data are representative of paired cultures from three different donors. C) Immunofluorescence using the antibodies against the indicated markers in three different primary human prostate explant cultures that clustered in the independent explant cohort (A) into C1/C2/C3 subpopulations. Font color denotes pseudo-coloring in the displayed merged images. Nuclei were counterstained using Hoechst 33342. Images are representative of three experiments using explant cultures from seven different patients.

**Supplemental Figure 5 Flow cytometry-based single cell analysis of primary human prostate fibroblast cultures**

A-H) Flow cytometry analysis of thirty-six biobank cultures selected randomly or on the basis of their substate annotation per the original transcriptomic dataset (Supplemental Table S2) and/or qRT-PCR of substate-delineating markers (Supplemental Fig. 4A) and stained for the cell surface stromal markers FAP, PDPN, CD90/THY1, CD140b/PDGFRB, CD140a/PDGFRA, MCAM and CD105/ENG. A-B) tSNE dimensional reduction analysis of flow cytometry (FC) data from twenty-three primary prostate fibroblast cultures (training set) based on their expression of the aforementioned stromal makers. Sample counts are stratified to visualize A) the original 13 clusters identified via XShift or B) sample number. C) Heatmap of flow cytometry data displaying percentage of counts in the positive marker gate and percentage of counts in samples annotated as C1, C2 or C3 subpopulations as defined and marked * in panel H). D) Gating strategy developed using the most distinguishable stromal surface markers identified from C), which yielded eight gated clusters from the original 13 clusters. E) tSNE plot overlaid with the 8 gated clusters identified by applying the gating strategy in D). F) Heatmap clustering based on the percentage of gated clusters for 36 fibroblast cultures (extended set), including the 23 explant culture samples from the training set. FC sample numbering of the extended set corresponds to that of the training set in the tSNE plot in A). Annotation of donor-related cultures that were also used in the qRT-PCR cohort (Supplemental Fig. 4A) and original transcriptomic cohort (Fig. 1, Supplemental Table 2) are depicted underneath the corresponding FC sample number. Data are scaled by column to highlight the most prominent cluster in each sample. G) tSNE plot overlaid with sample counts from cultures annotated as C1, C2 or C3 subpopulations as defined and marked * in panel H). Grey shading denotes ungated cells of samples that did not display a single enriched cluster or were not determined (n.d.). H) Stacked bar plot displaying the percentages of gated clusters within each primary prostate fibroblast culture sample (n = 36), grouped according to the predominant gated cluster composition of each sample. Sample numbering beneath each bar corresponds to the sample numbering in F). Cultures enriched for a single cluster (≥55%) were annotated as C1, C2 or C3 (marked *). Cultures enriched for two of either clusters 6 and 10 or clusters 10 and 1 were annotated as C1/C2 or C2/C3, respectively. All other cultures were designated n.d. (not determined). I) qRT-PCR of the indicated gene in primary prostate fibroblast cultures sorted via FACS for cluster 1, 6+10 combined (*lower panel)* or cluster 6 or 10 separately (*upper panel)*. Genes associated with general fibroblast activation/contractile markers are indicated in black, while C1, C2, and C3 subpopulation-delineating markers (from Fig. 1) are shown in blue, yellow, and red, respectively. Values represent mean $2^{-\Delta\Delta Ct}$ via qRT-PCR, normalized to the housekeeping gene TBP. Data are representative of two independent sorting experiments using five fibroblast cultures isolated from different donors. F-H) Font color denotes the subpopulation annotated using the different complementary methodologies (C1, blue; C2, yellow; C3, red; black font, not determined; black font with yellow-red box denotes cultures considered to comprise a mix of C2/C3 substates).

**Supplemental Figure 6 Multiplex immunofluorescent and immunohistochemical staining of human PCa reveals distinct CAF subpopulations *in vivo* – pertaining to Figure 2**

A-C) Human prostate tissue sections of indicated pathology were stained using the antibodies shown. Original magnification 200x. Font color denotes pseudo-coloring in the displayed images. A-B) Nuclei were counterstained using Hoechst 33342 (blue). Boxed regions are shown enlarged beneath each parental image. B) Four-channel merged image of a negative control incubated without primary antibodies. C) Immunohistochemistry of consecutively stained sections of indicated pathology using the antibodies indicated. D) Immunohistochemistry of CES1 in low-grade PCa. Boxed region is enlarged right. A-C) Images are representative of 4 independent experiments using tissue sections derived from 8 different patients. E-G) Expression of SMC signature genes (combined z-score) in the TCGA-PRAD cohort using the indicated signatures. H-I) Kaplan-Meier plots depicting progression-free survival (PFS) of high-grade (≥T stage T2c) samples of the TCGA-PRAD cohort stratified as described in Materials and Methods for high ($SMC^{high}$) or low ($SMC^{low}$) expression levels of the indicated SMC signatures. Source data for panels C-I are provided in the Source Data file (Supplemental Table 6).

**Supplemental Figure 7 AR loss in myofibroblastic CAF renders them insensitive to the proliferative effects of enzalutamide - pertaining to Figure 4**

A) Western blotting of thirteen partially patient-matched explant cultures using the antibodies indicated (original western blot sourcing the densitometric data shown in Fig. 5D). B) Densitometric-based correlation of AR and pSTAT3 or ITGA11 protein levels in thirteen explant cultures as depicted in A). C-D) Immunofluorescent staining of human prostate cancer tissue sections using the antibodies indicated. Font color denotes pseudo-coloring in the displayed merged images. Nuclei were counterstained using Hoechst 33342. Boxed regions are shown enlarged, right. Arrowheads in panel 2 indicate non-SMC ($CCDC102B^-$) $AR^+$ stromal cells. E-F) ImageJ quantification of tissues stained as in D) whereby the area stained by the indicated antibody is expressed as a percentage of the total $DAPI^+$ area, panel E). For AR colocalization in panel F) the percentage area is shown for $AR^+$ nuclei ($DAPI^+$ and $AR^+$) also positive for each stromal cell type marker (e.g. $DAPI^+$ and $AR^+$ and $ENG^+$). The quantification does not distinguish between $ENG^+$ myCAF and $ENG^+$ endothelial cells, the latter comprising a key source of $ENG^+AR^+$ cells in high grade stromogenic tumors. Data represent mean ± SEM of at ten fields of view from three different high-grade tumors stratified according to their stromogenic or non-stromogenic content defined as myCAF-rich ($ENG^+$/$ITGA11^+$) or SMC-rich ($SMA^+$/$CNN1^+$). G) qRT-PCR of *FKBP5* (left) and *FAP* (right) in C1 cultures in steroid hormone-depleted medium treated with or without 10 nM R1881 in the presence of 10 µM enzalutamide (ENZA) or vehicle equivalent for 72 h.

H-I) Western blotting of the indicated explant culture under hormone-replete conditions in the presence of 10 µM enzalutamide or vehicle equivalent for 72 h. H) Values denote densitometric intensity relative to the corresponding control treated culture. I) Source western blot depicted in Fig. 4K. J) Densitometric quantification of pAKT levels in fibroblast cultures incubated for 72h with 10 µM enzalutamide (ENZA) or vehicle equivalent (related to Fig. 4L). K-M) Real time proliferation of the indicated fibroblast cultures incubated K) in 10% steroid hormone-replete DMEM with 10 µM enzalutamide or vehicle equivalent (ctrl) or L-M) in 2.5% steroid hormone-deplete DMEM with the indicated concentration of R1881 and 10 µM enzalutamide or vehicle equivalent (related to Fig. 4M-P). N-O) Migration assay of C1 fibroblasts pretreated for 96h with the indicated concentration of R1881 and 10 µM enzalutamide or vehicle equivalent (ctrl). Statistical significance was calculated: B) Pearson correlation, E-F) two-way ANOVA with Tukey's multiple comparison correction, G) one-way ANOVA with Holm-Šídák multiple comparison correction, J) Mann-Whitney test, K-N) two-way ANOVA with Holm-Šídák multiple comparison correction. C, D, H, I, O) images are representative of at least three independent experiments using primary material from different donors.

**Supplemental Figure 8 Dynamic remodeling of CAF activation states *in vivo* - pertaining to Figure 5**

A) Sample level heatmaps depicting expression level of the corresponding murine orthologs for the indicated explant culture signature genes (related to Fig. 5B) in the mouse transcriptome of castrated and intact samples from the indicated PDX model. B-C) Single channel monochromatic images of images displayed in Fig. 5D and Supplemental Fig. 7D of the indicated PDX tumor using the antibody indicated. D) Additional fields of view of the indicated PDX tumor stained using the antibodies indicated. E) Immunohistochemistry of mouse Itga11 or IgG control in LAPC9 intact tumors. Enlarged images of boxed regions are shown beneath each parental image. (B-E) Images are representative of at least two independent experiments using tissue sections from at least two different tumors per condition.

**Supplemental Figure 9 NFκB-YAP-TGFβ signaling axis underlies AR loss during fibroblast phenotypic switching - pertaining to Figure 6**

A) qRT-PCR of C1 and C3 cultures treated with 1 µM SB431542 or vehicle control for 72h. Values represent mean fold change ± SEM in expression relative to vehicle control treated cells. B) Densitometric quantification of C3 cultures treated as in A). Related to Fig. 6C. C) Morphology and D) mean fold change gene expression (qRT-PCR) of the indicated fibroblast cultures grown as 3D spheroids relative to standard 2D conditions for 4 days. E) Morphology and F) mean fold change gene expression (qRT-PCR) of two C3 cultures on 2 kPa soft hydrogels relative to standard tissue culture plasticware (TCP) for 4 days. G) Sample level heatmap showing expression of YAP1 target genes in the explant transcriptomic dataset. H-J) Brightfield imaging of three C3 cultures isolated from independent patients and treated with the indicated compound for 96 h (10 nM R1881, 1 µM SB431542, 10 µM enzalutamide (ENZA), 1 µM verteporfin, 2 µM BAY11-7082 or vehicle equivalents). Related to Fig. 6E and 6G. Scalebars denote 200 µm. Statistical significance was calculated: A) two-way ANOVA with Šídák multiple comparison correction, B) Wilcoxon signed rank test, E) Mann Whitney U test, G) one-way ANOVA using Tukey multiple comparison correction. C, E, H-J) images are representative of at least three independent experiments using primary material from different donors.

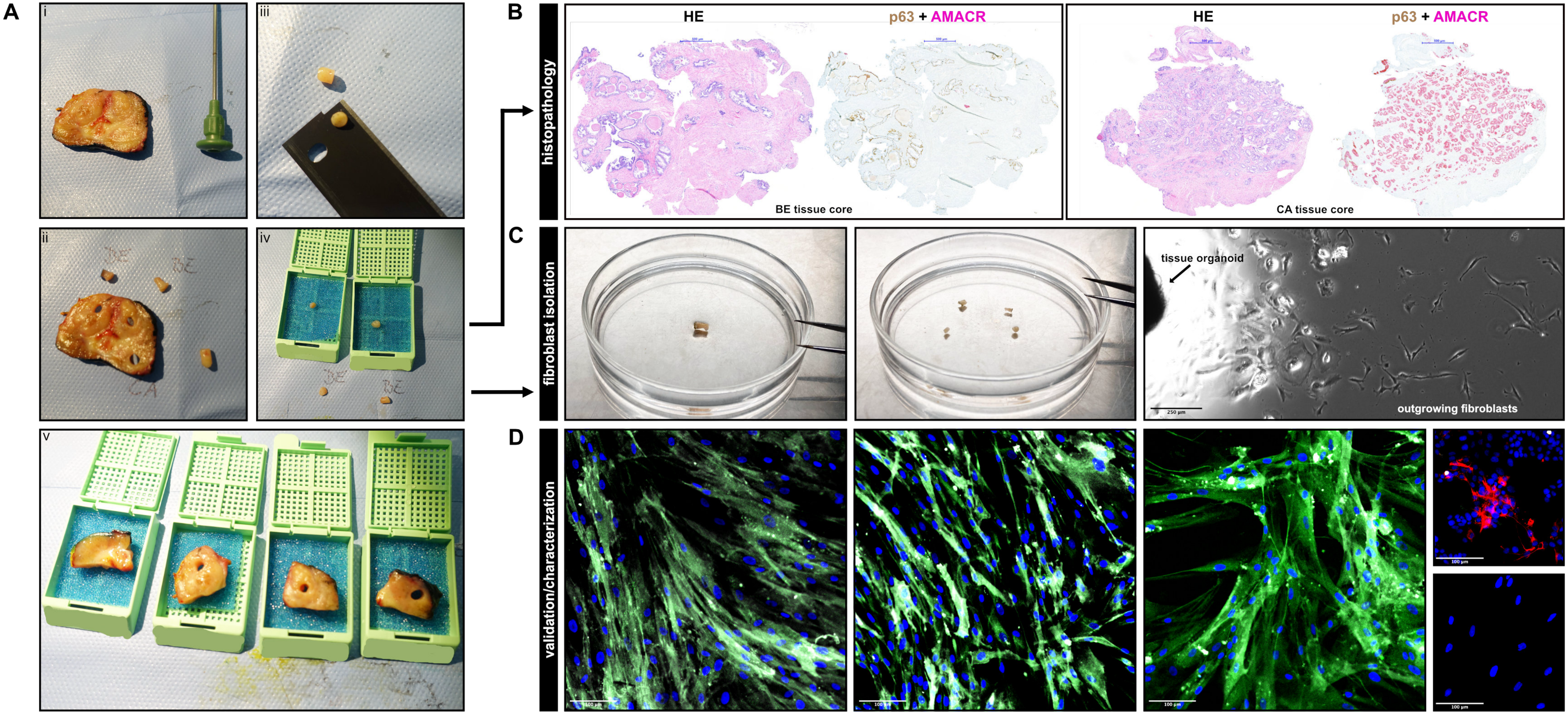


Supplemental Fig. 1

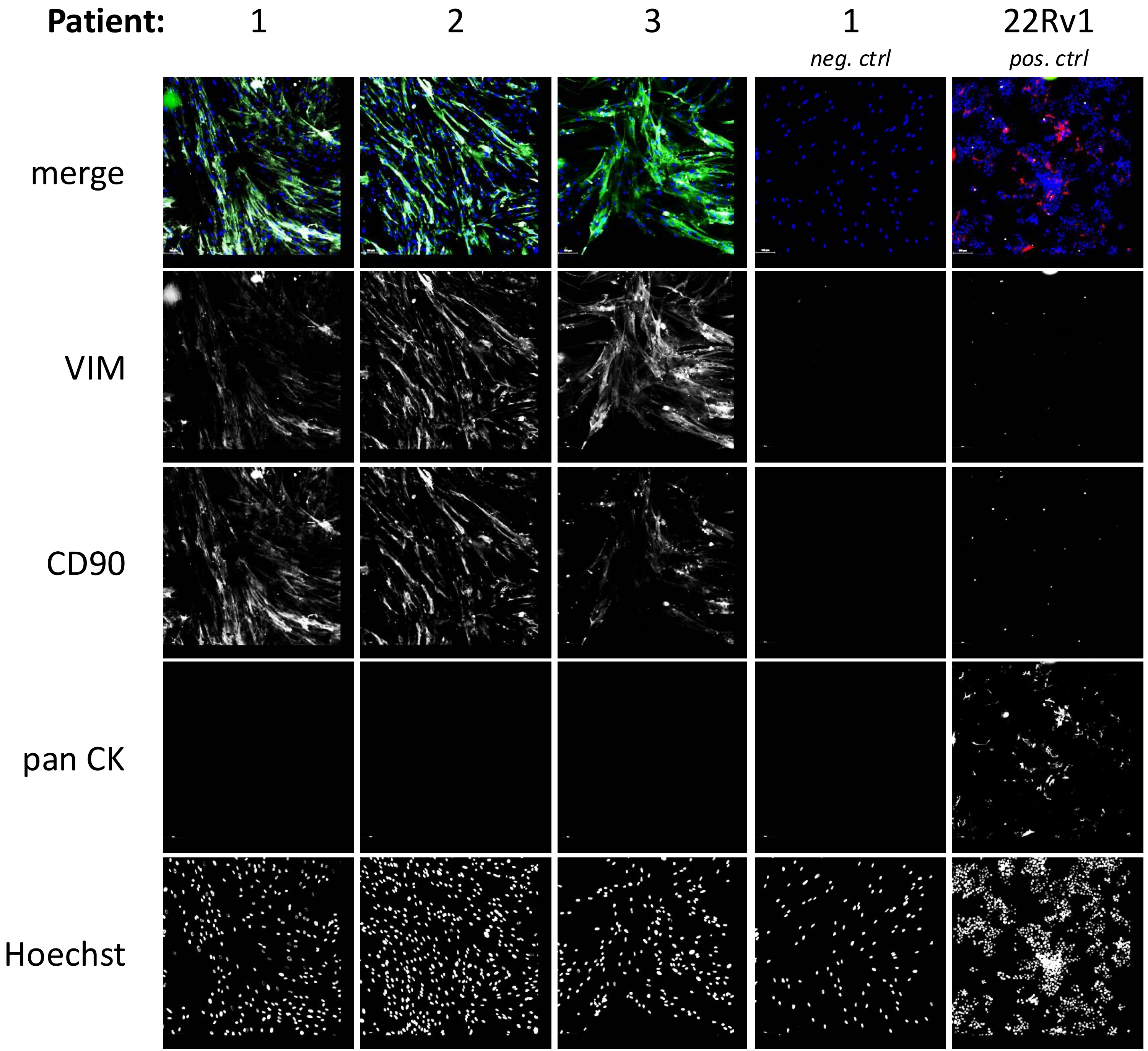


**Supplemental Fig. 2**

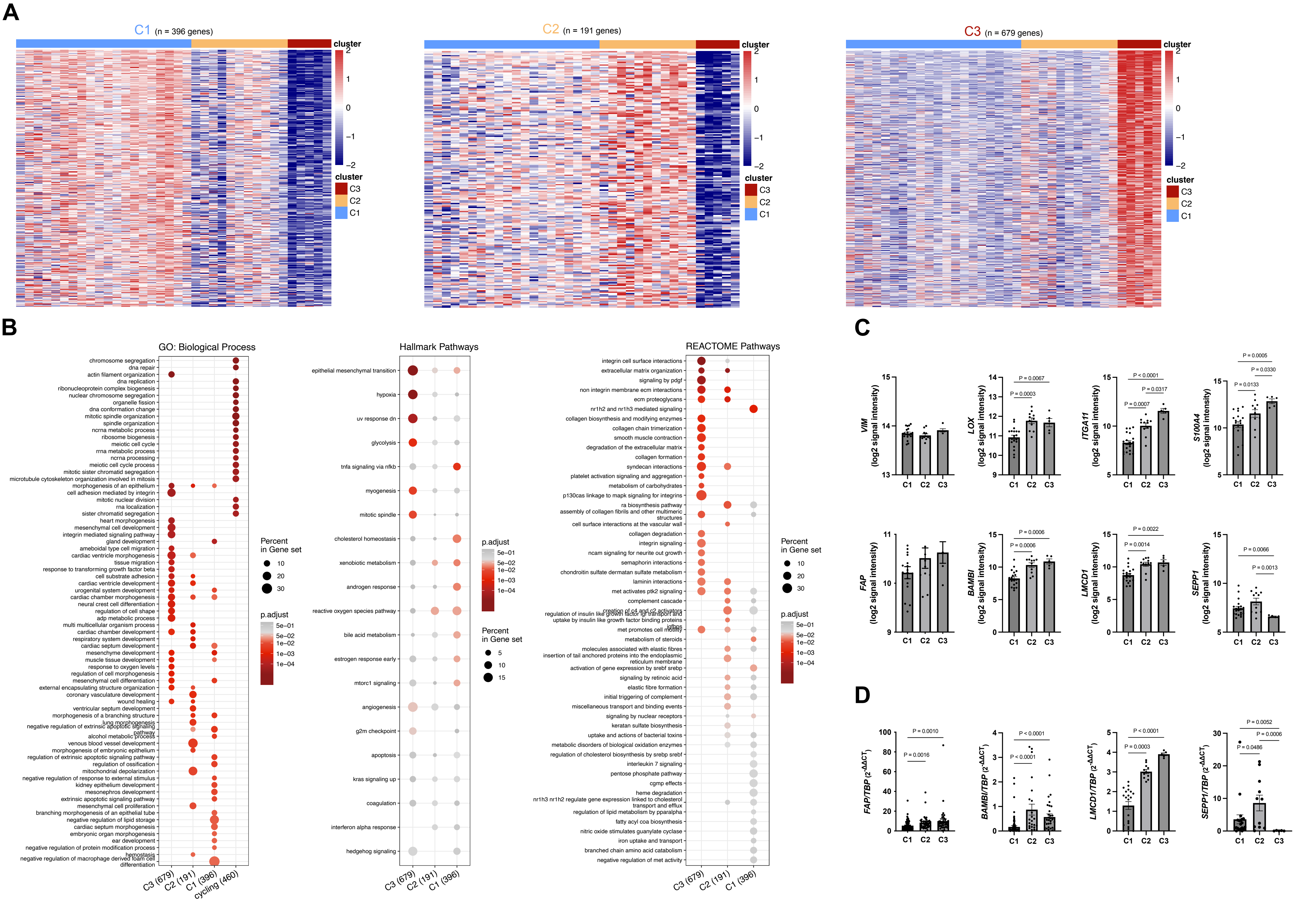


Supplemental Fig. 3

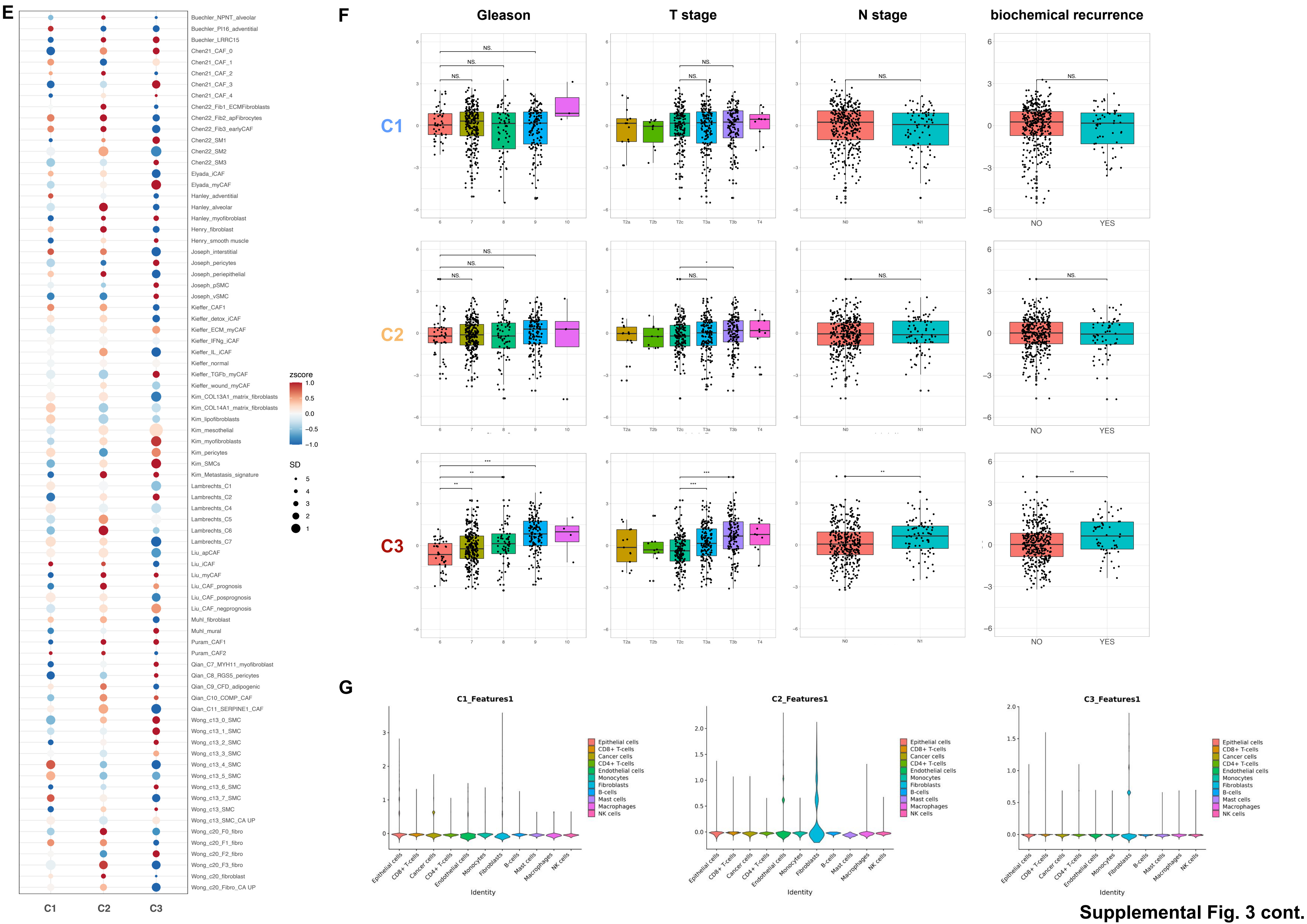


Supplemental Fig. 3 cont.

Supplemental Fig. 4

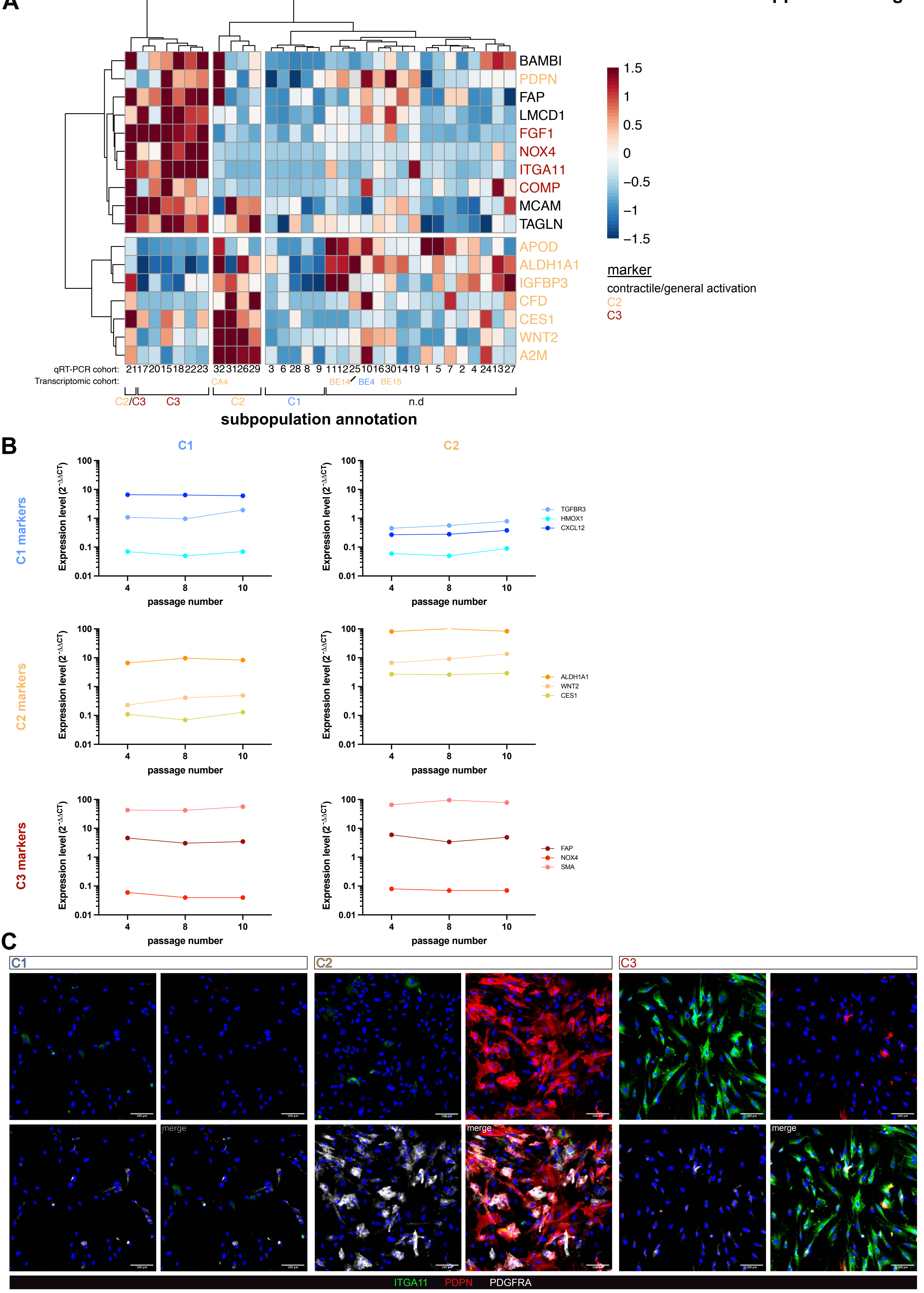

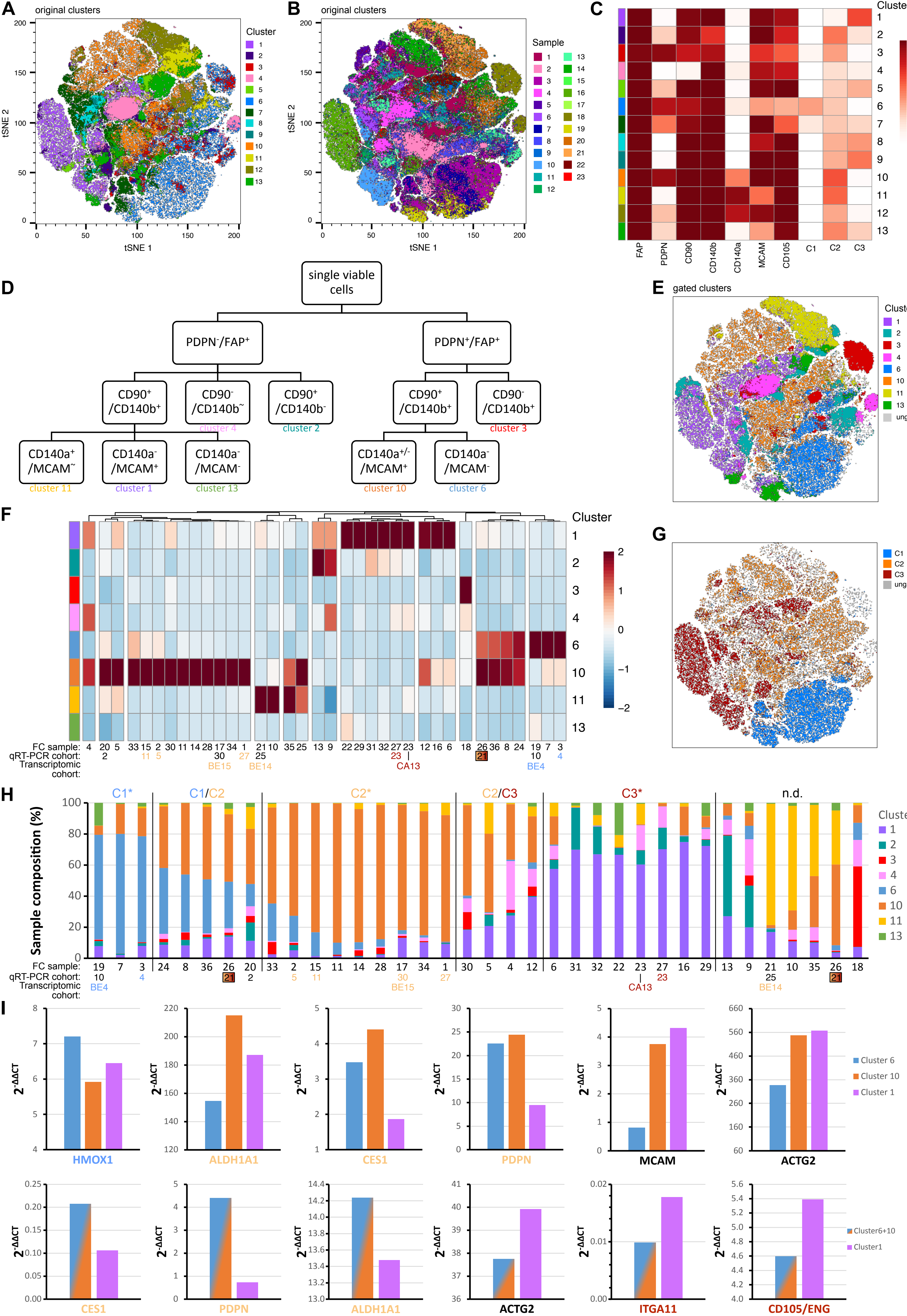


Supplemental Fig. 5

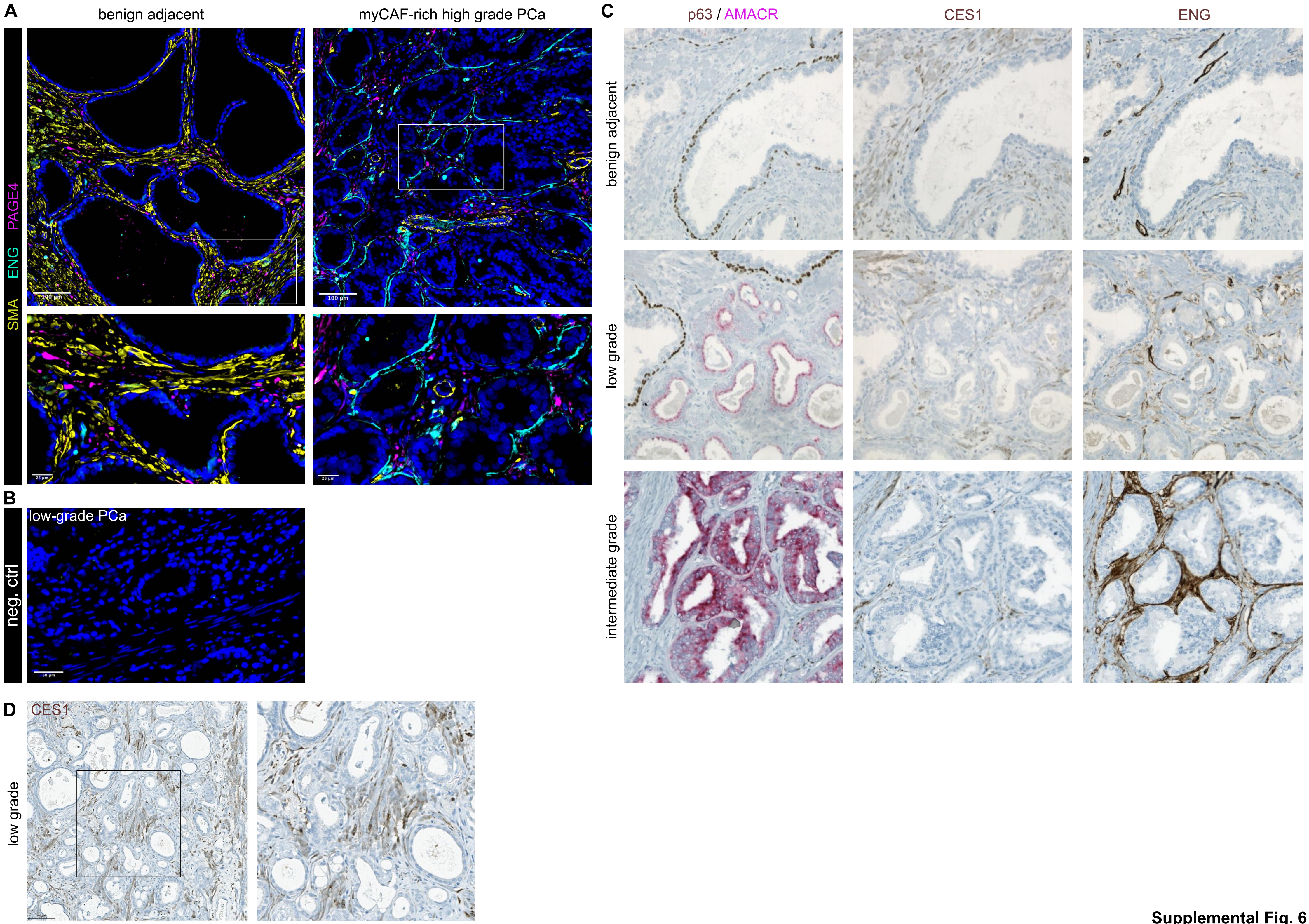


Supplemental Fig. 6

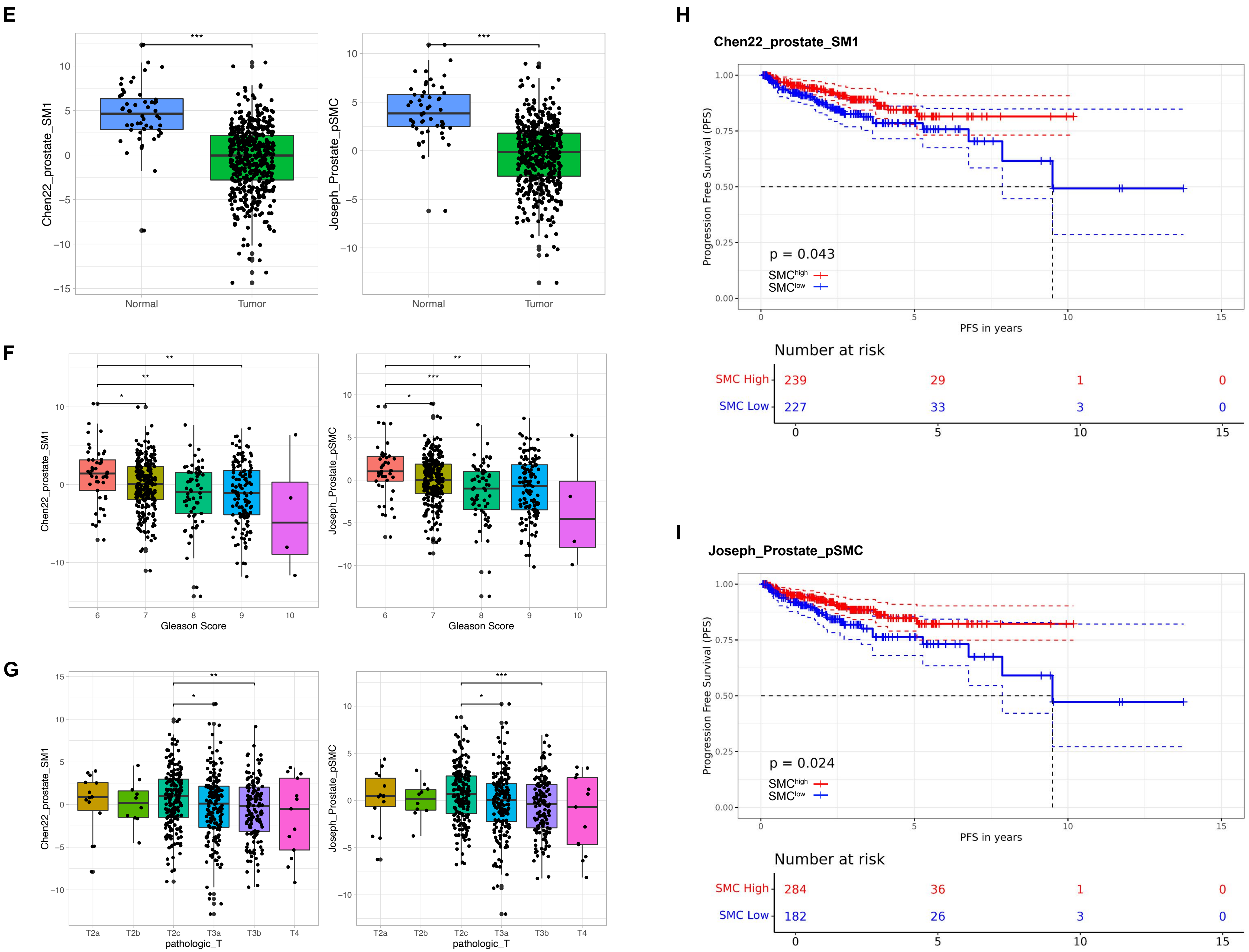


**Supplemental Fig. 6 cont.**

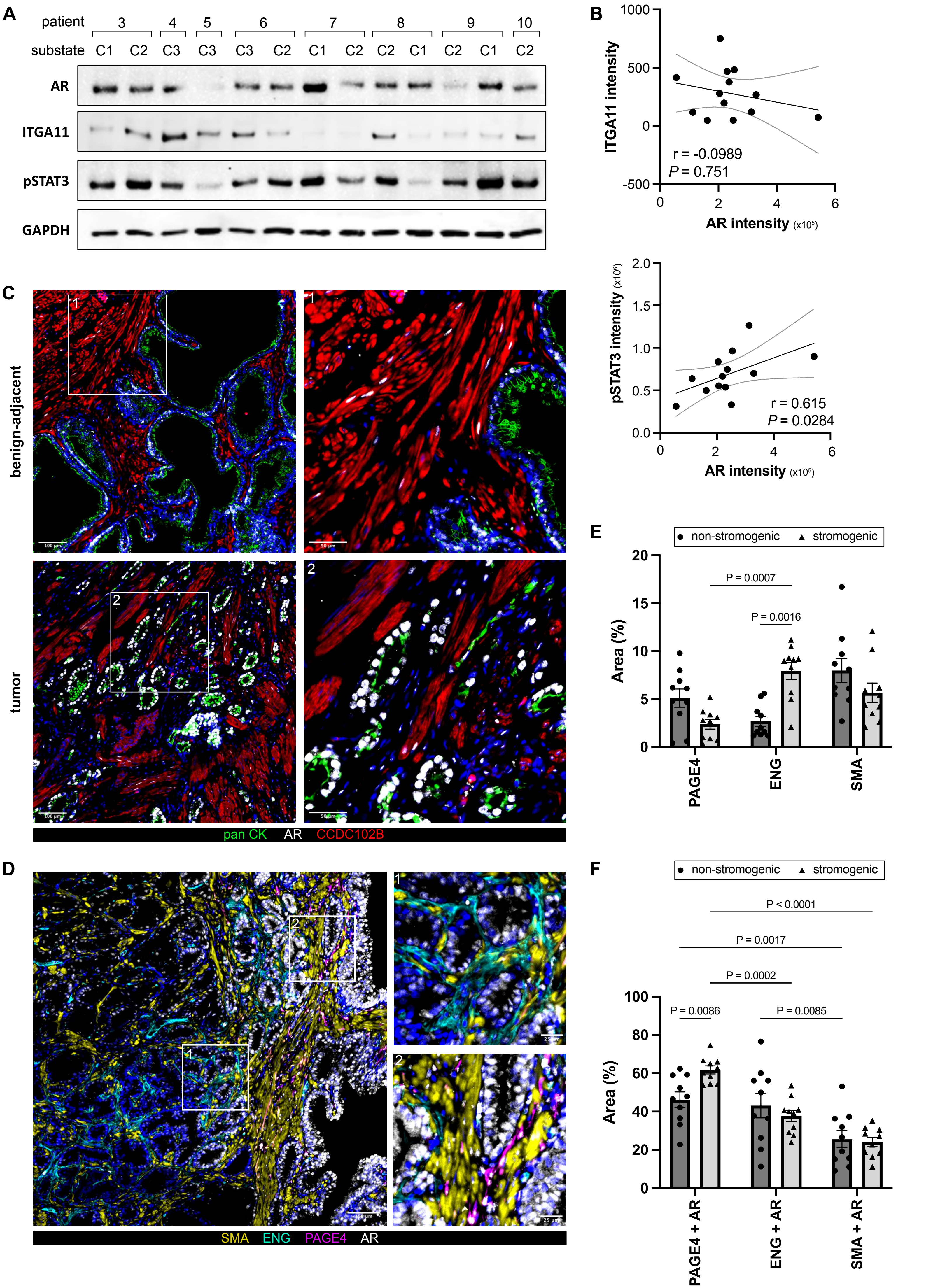
A
patient 3 4 5 6 7 8 9 10
substate C1 C2 C3 C3 C3 C2 C1 C2 C2 C1 C2 C1 C2
AR
ITGA11
pSTAT3
GAPDH
B
ITGA11 intensity
AR intensity (x10^5)
r = -0.0989
P = 0.751
pSTAT3 intensity (x10^6)
AR intensity (x10^5)
r = 0.615
P = 0.0284
C
benign-adjacent
tumor
pan CK AR CCDC102B
D
SMA ENG PAGE4 AR
E
non-stromogenic stromogenic
Area (%)
P = 0.0007
P = 0.0016
PAGE4
ENG
SMA
F
non-stromogenic stromogenic
Area (%)
P < 0.0001
P = 0.0017
P = 0.0002
P = 0.0086
P = 0.0085
PAGE4 + AR
ENG + AR
SMA + AR

Supp. Fig. 7

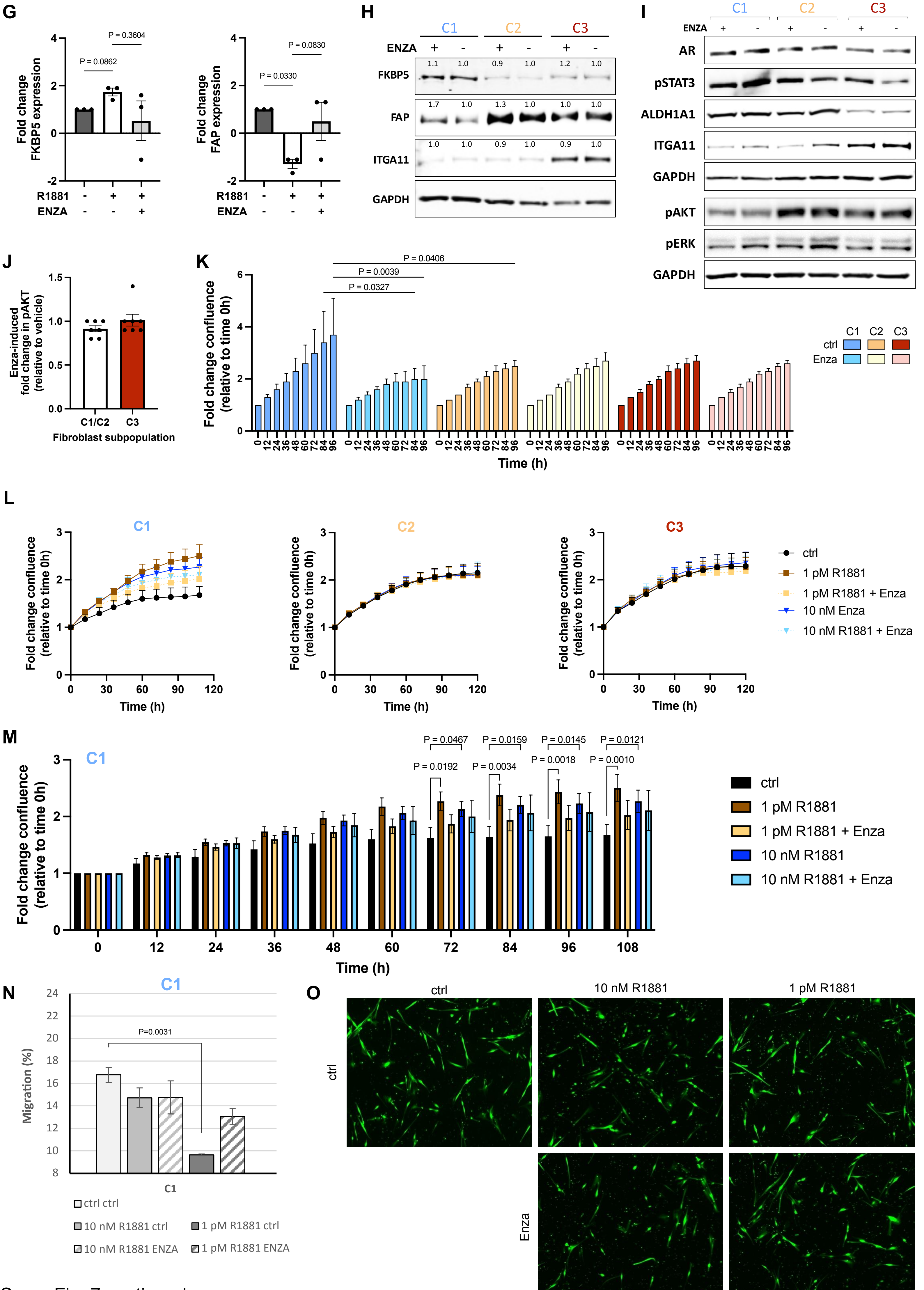


Supp. Fig. 7 continued

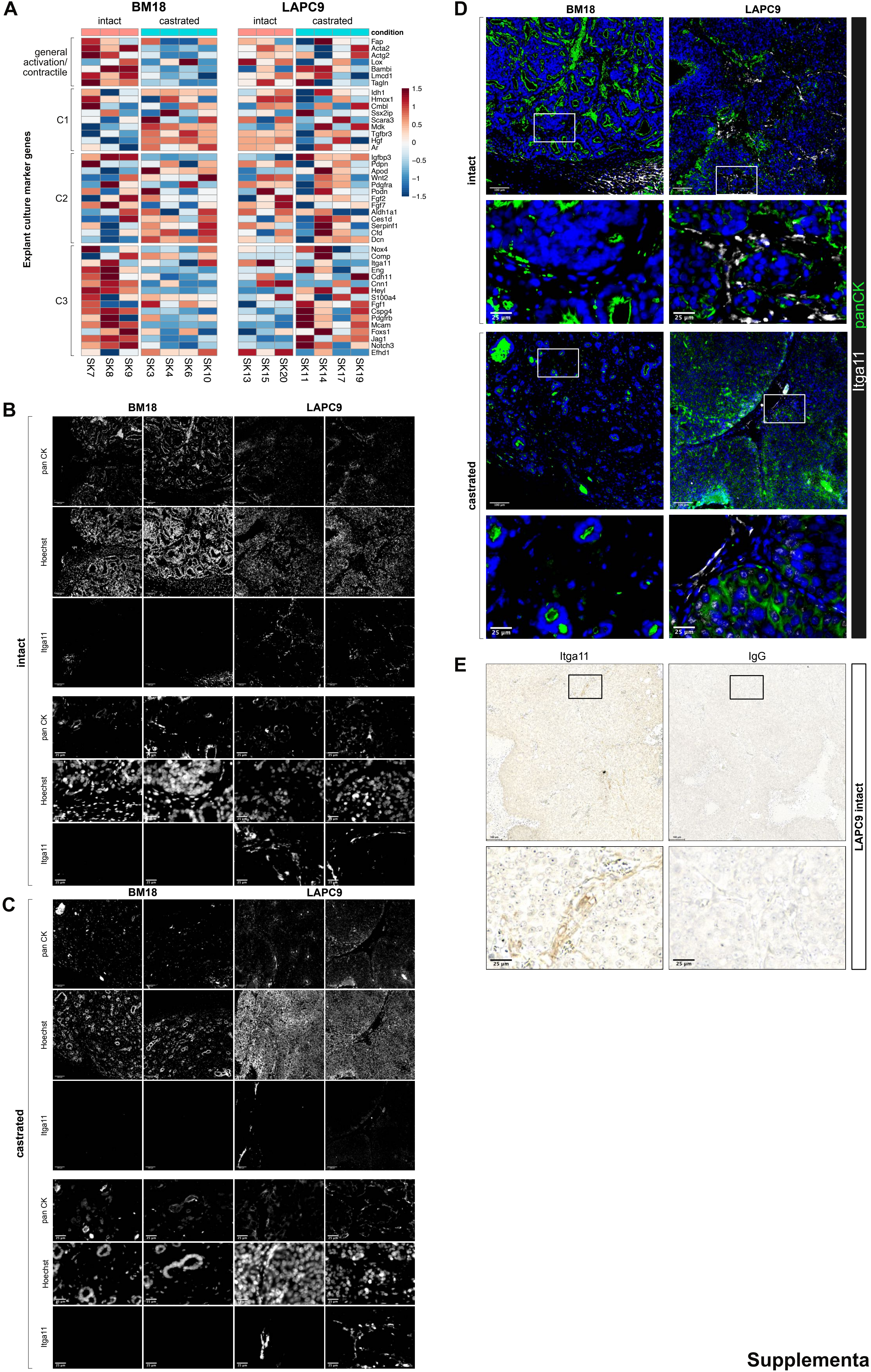


Supplemental Fig. 8

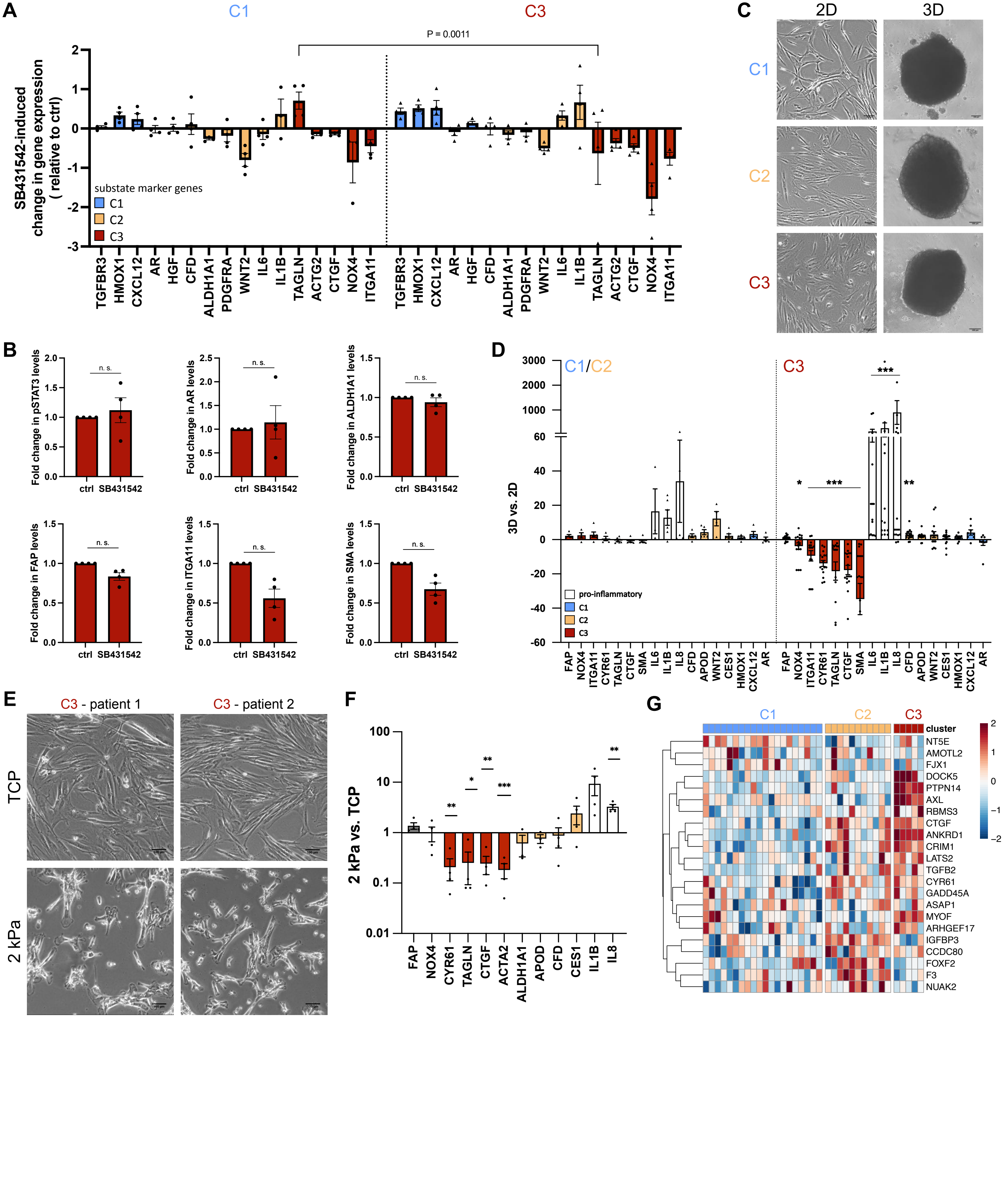


Supplemental Fig. 9

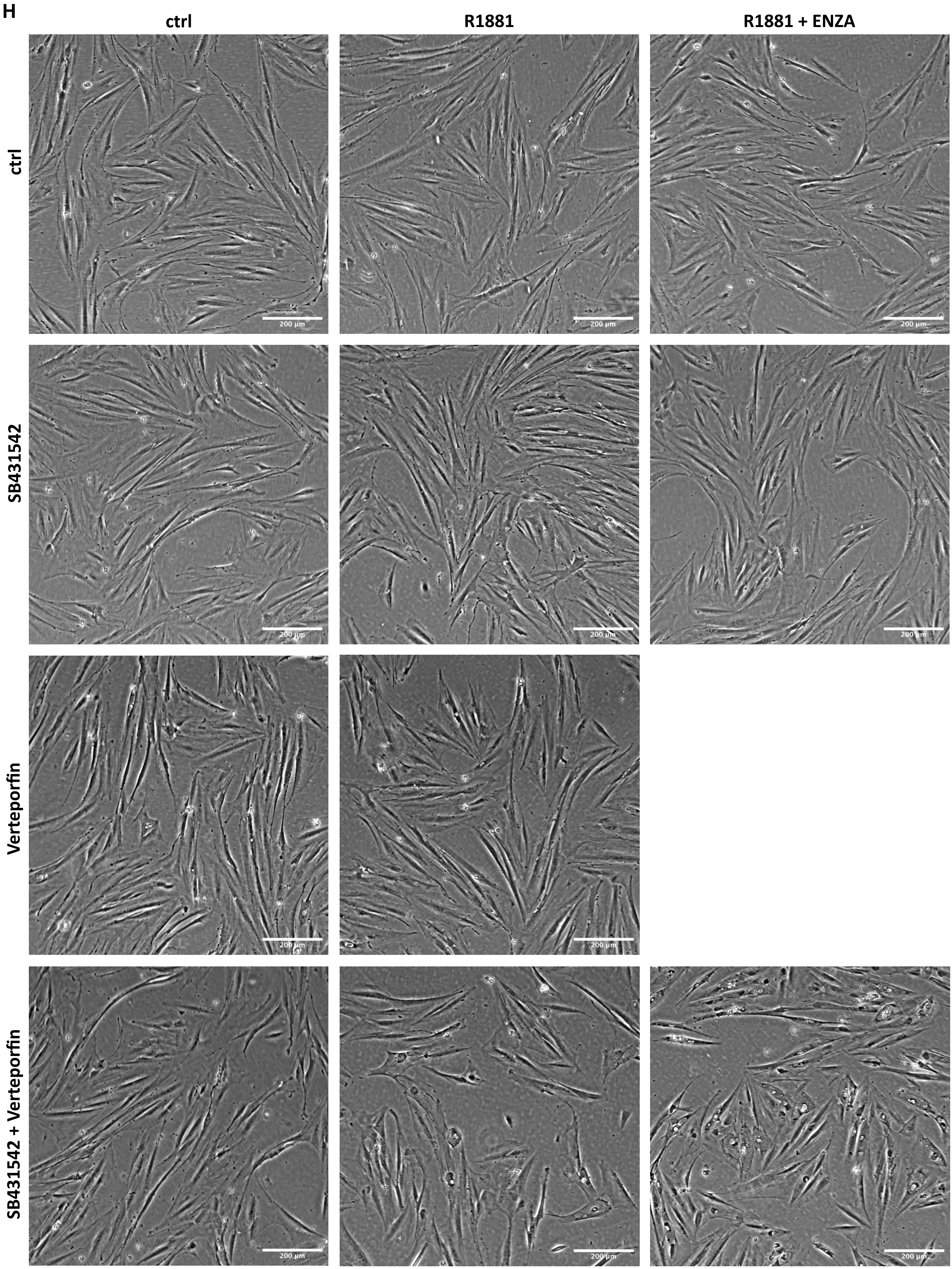
H
ctrl
R1881
R1881 + ENZA
ctrl
SB431542
Verteporfin
SB431542 + Verteporfin
200 µm

I

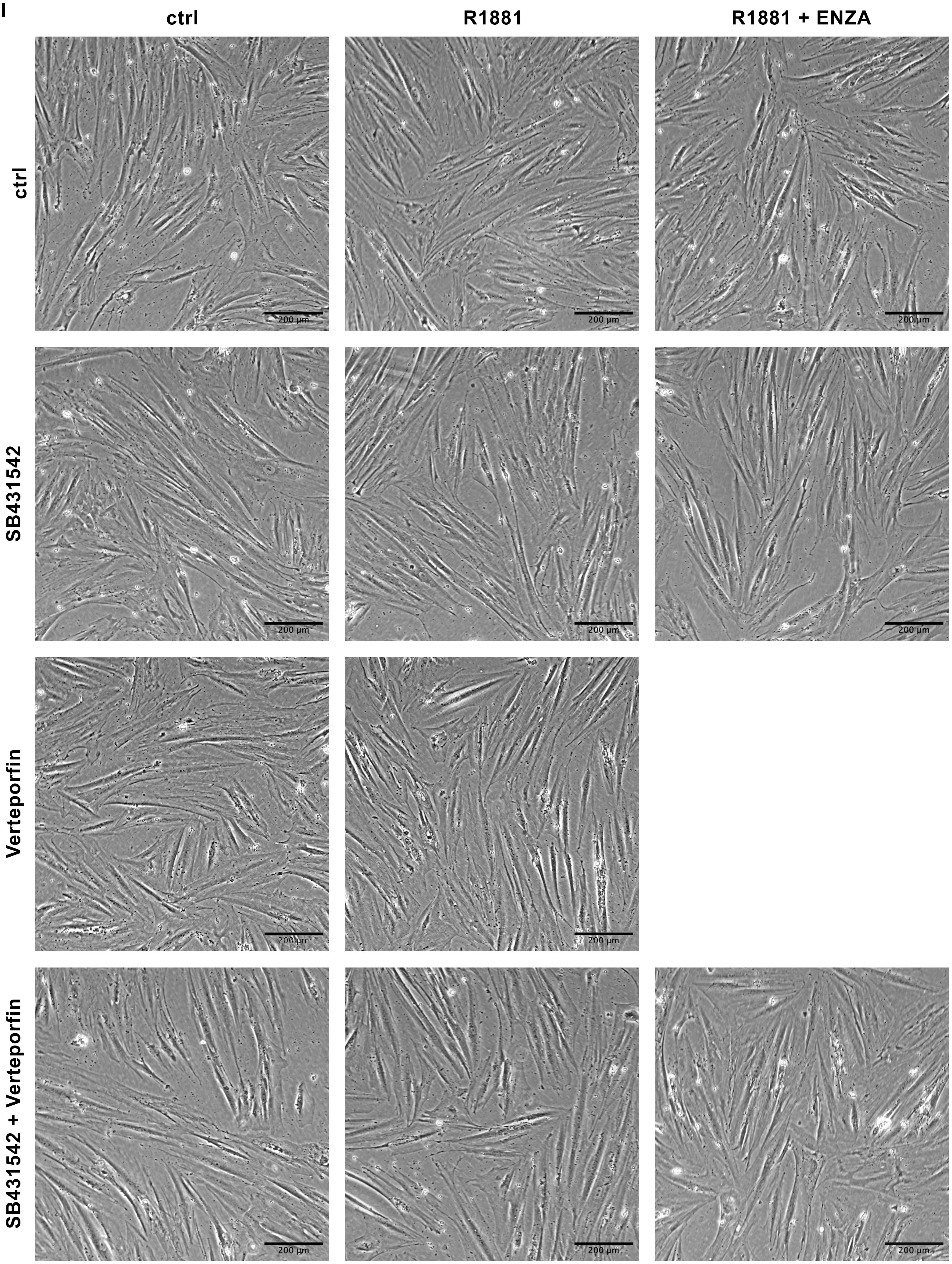


Supplemental Fig. 9 cont.

I (continued)

ctrl | BAY11-7082 | BAY11-7082 + Verteporfin

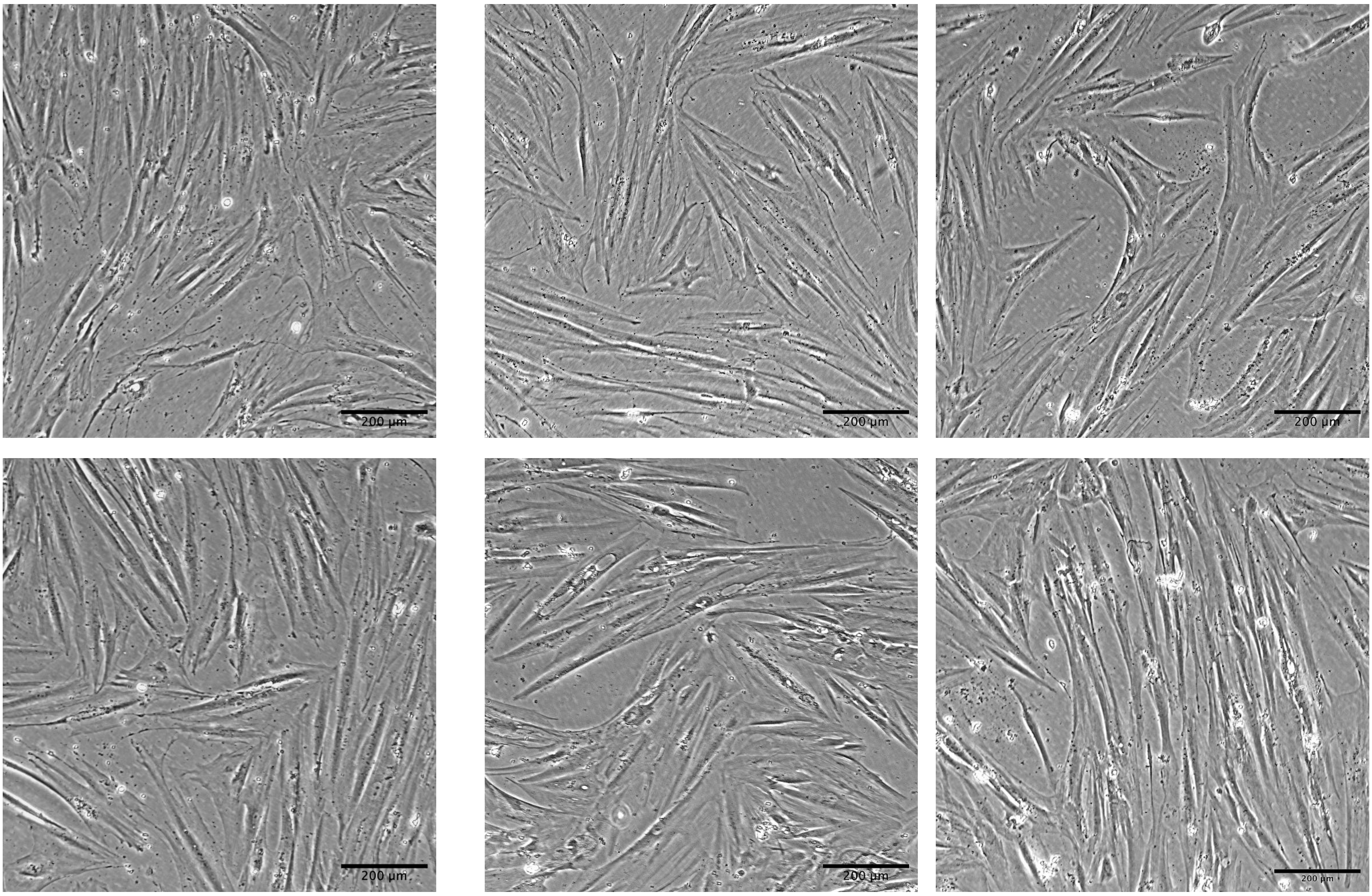


Verteporfin + SB431542 + R1881 | BAY11-7082 + Verteporfin + R1881 | BAY11-7082 + Verteporfin + R1881 + SB431542

Supplemental Fig. 9 cont.

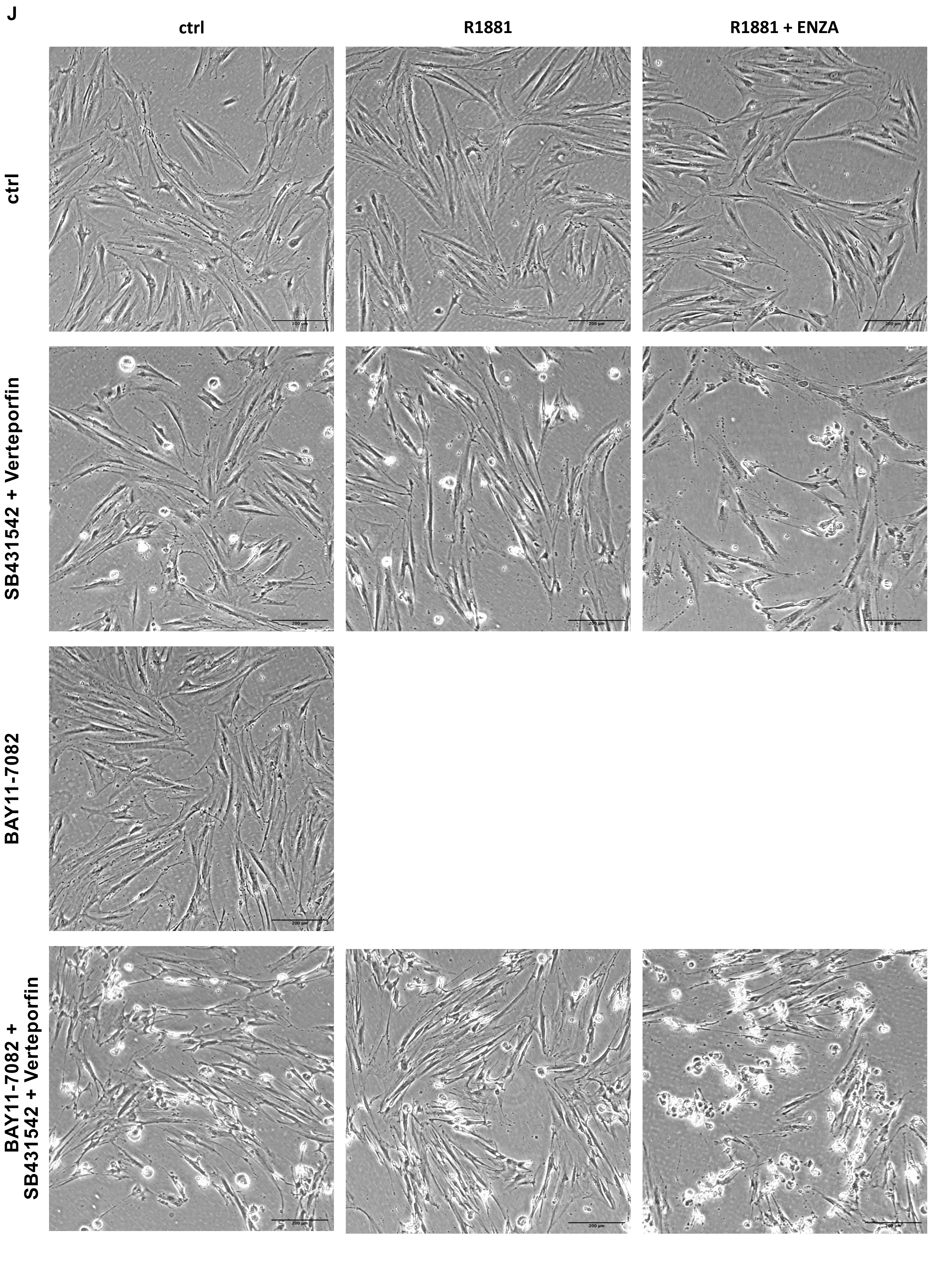


Supplemental Fig. 9 cont.
